# An endogenous cluster of target-directed microRNA degradation sites induces decay of distinct microRNA families

**DOI:** 10.1101/2024.12.11.627053

**Authors:** Nicholas M. Hiers, Lu Li, Tianqi Li, Peike Sheng, Yuzhi Wang, Conner M. Traugot, Michael Yao, Mingyi Xie

## Abstract

While much is known about miRNA biogenesis and canonical miRNA targeting, relatively less is understood about miRNA decay. The major miRNA decay pathway in metazoans is mediated through target-directed miRNA degradation (TDMD), in which certain RNAs can “trigger” miRNA decay. All known triggers for TDMD base pair with the miRNA seed, and extensively base pair on the miRNA 3′ end, a pattern that supposedly induces a TDMD-competent conformational change of Argonaute (Ago), allowing for miRNA turnover. Here, we utilized Ago1-CLASH to find that the *Drosophila* transcript *Kah* contains at least two triggers, a “trigger cluster”, against miR-9b and the miR-279 family. One of these triggers contains minimal/non-canonical 3′ end base pairing but is still sufficient to induce TDMD of the entire miR-279 family. We found that these clustered triggers likely lack cooperativity, the minimal 3′ pairing is required for miR-279 family turnover, and probed the in-cell RNA structure of the *Kah* trigger cluster. Overall, this study expands the list of endogenous triggers and the unexpectedly complex regulatory network governing miRNA degradation.

## INTRODUCTION

MicroRNAs (miRNAs) are a class of small ∼22 nucleotide (nt) non-coding RNAs that induce post-transcriptional gene silencing^1^. To do this, miRNAs are bound by one of the Argonaute (AGO) family proteins and serve as guides for AGO association with target RNAs^1–3^. Typically, a target bearing sequence complementarity to miRNA seed (nt 2-8) is sufficient to predict down-regulation^4,5^. With such a short regulatory sequence requirement, and hundreds of conserved miRNA genes across metazoans, it is unsurprising that miRNAs are thought to act as “master” regulators of post-transcriptional gene expression. Indeed, loss of individual miRNA genes has been shown to induce a variety of phenotypes including developmental abnormalities and embryonic lethality^1,6,7^.

While extensive research has been given to the study of miRNA biogenesis and functional consequences, relatively less is understood about miRNA decay^8^. For many years, researchers had observed that certain miRNA targets bearing extensive 3′ complementarity (in addition to seed-matching) could “trigger” rapid miRNA turnover, a process collectively referred to as target-directed miRNA degradation (TDMD)^9–15^. In 2020, it was revealed that the endogenous TDMD mechanism in mammals is catalyzed via ZSWIM8, a Culin-RING E3 ubiquitin ligase^16,17^. Mechanistically, it appears that when miRNAs bind to TDMD targets, hereafter simply referred to as TDMD “triggers”, the extensive base pairing induces a conformational change in AGO, enabling ZSWIM8 recognition and AGO polyubiquitination^16–19^. This process directs AGO for proteasomal decay, in effect exposing the miRNA to cellular ribonucleases upon loss of AGO. Since this revelation, several studies have identified the ZSWIM8 ortholog in *Drosophila* (Dora) and *C. elegans* (Ebax-1) also carry out TDMD in their respective systems^17,20–23^. In fact, several recent reports suggest TDMD plays a fundamental role in both mammalian and *Drosophila* development^21,24,25^. For example, TDMD of the miR-3 family is required for proper *Drosophila* embryogenesis, and the loss of such degradation appears lethal *in vivo*^21^. Also, in mammals, it was recently reported that TDMD of miR-322/503 is required for proper mammalian growth^24^. In total, loss of ZSWIM8 (or Dora/Ebax-1) increases the abundance of nearly 100 miRNAs; however, the triggers that induce TDMD for most of these miRNAs are still unknown^17,20–22,24–26^. Therefore, it is essential to identify each of these triggers to better understand the TDMD molecular mechanism and phenotypic consequences of TDMD in metazoans.

There have been two main methods used to successfully screen for and validate novel triggers: computational algorithmic screens, and biochemical methods-based approaches^21,27–30^. In either case, these studies typically employ a stringent extensive 3′ complementarity requirement for putative triggers, given that all known TDMD examples contain at least 7 consecutive base-pairings between triggers and the miRNA 3′ end^26,31^. Interestingly, it was recently revealed that there is likely a “seed-sufficient” trigger in *C. elegans* that requires no 3′ end complementarity to degrade the miR-35 family^20^. Relatedly, extensive target complementarity with miRNAs can out-compete the miRNA 3′ association with the AGO PAZ domain, thereby exposing the miRNA 3′ end to non-templated nucleotide addition by terminal nucleotidyltransferases (TENTs)^10–13,18,19,32,33^. Given that all known triggers extensively base pair with their targets, they have also been reported to induce tailing/trimming of associated miRNAs^18^. However, it was recently observed that several examples of miRNAs are stabilized following loss of ZSWIM8 (ZSWIM8-sensitive) without any change in tailing^24^. Together these observations suggest that there are potential triggers in metazoans that base pair less extensively with the cognate miRNA but are still sufficient to induce TDMD.

We recently have had success in the application of AGO-crosslinking, ligation, and sequencing of hybrids (AGO-CLASH)^34^, to screen for TDMD triggers^29,30^. AGO-CLASH is a modified CLIP method which utilizes UV crosslinked cells, immunoprecipitation of AGO, and an intermolecular ligation between the miRNA and target RNA^34–37^. Sequencing of the subsequent miRNA-target hybrid enables identification of high-confidence miRNA-target interactions. Our application of this method in *Drosophila* S2 cells allowed for the identification of 5 triggers: *Ago1*:miR-999, *h*:miR-7, *Kah*:miR-9b, *Wgn*:miR-190, and *Zfh1*:miR-12^30^. Given that many other miRNAs are stabilized following loss of Dora in S2 cells, there are additional triggers remaining that require identification^17,21,23^.

In the initial studies, we employed stringent extensive base pairing criteria for the screening of potential triggers^29,30^, though AGO-CLASH data could presumably be used to identify even non-canonical triggers by simply relaxing the screening criteria. Here, we find that the *Kah* transcript, which we originally identified as a miR-9b trigger, contains a secondary trigger sufficient to induce decay of the entire miR-279 family, consisting of miR-279, miR-286 and miR-996. Interestingly, this quality makes *Kah* an endogenous example of multiple triggers within the same transcript, a “trigger cluster”. Surprisingly, this miR-279 family trigger contains non-canonical TDMD base pairing, in that relatively little 3′ complementarity is present, though this minimal 3′ complementarity still appears required for miR-279 family turnover. In addition, in-cell structural probing of the endogenous *Kah* trigger cluster suggests the seed-binding regions of these triggers reside in accessible single-stranded regions, whereas the 3ʹ end binding regions appear more structured. Our study suggests that the types of base pairs that induce TDMD are far from full comprehension. Overall, our results shed light on the existence of both non-canonical and clustered triggers and highlight the ability of AGO-CLASH to aid in identifying additional unexpected triggers.

## RESULTS

### Ago1-CLASH suggests a cluster of TDMD triggers in *Kah*

In our initial screen for triggers in *Drosophila* S2 cells, we validated each of our highest confidence candidates via CRISPR-Cas9-mediated deletion^30^. In all but one case, loss of the trigger increased the abundance of the predicted miRNA specifically. With knockout (KO) of the *Kah* miR-9b trigger (*Kah*-9b), there is a significant increase in the abundance of both the miR-9b and miR-996 guide strands but not their co-transcribed passenger strands (Figures 1A and 1B). This quality is crucial to differentiate miRNA increase in abundance due to loss of TDMD (increase guide abundance), from an increase in biogenesis (increase both guide and passenger abundance)^12,17,20–22,24,25,29,30^. Puzzlingly, miR-9b and miR-996 belong to distinct miRNA families and therefore have different seed sequences and target repertoires: miR-9b is a member of the miR-9 family (miR-9a/b/c), while miR-996 is a member of the miR-279 family (miR-279/286/996). Upon further scrutiny, we observed that the *Kah*-9b trigger KOs broadly upregulated the guide strands of the whole miR-279 family (Figure 1A and 1B). Given the possibility of ligation biases that could disproportionally reflect miRNA abundance in our previously outsourced small RNA-seq^38–41^, we validated the upregulation of both miR-279 and 996 via near-infrared northern blot (from here on, northern blot)^42^. As in the *Dora*-KO, disrupting the *Kah*-9b trigger elevated the levels of miR-9b, miR-279 and miR-996, while the control miRNA bantam did not increase abundance (Figure 1C). It should be noted that miR-286 is likely undetectable by northern blot in S2 cells due to its relatively low abundance and therefore was excluded in this analysis. Since the miR-279 family guides are upregulated upon loss of Dora^17^ and are similarly stabilized following loss of *Kah*-9b (Figures 1C and S1A), we therefore considered whether we may have unintentionally perturbed a potential miR-279 family trigger.

**Figure 1.**
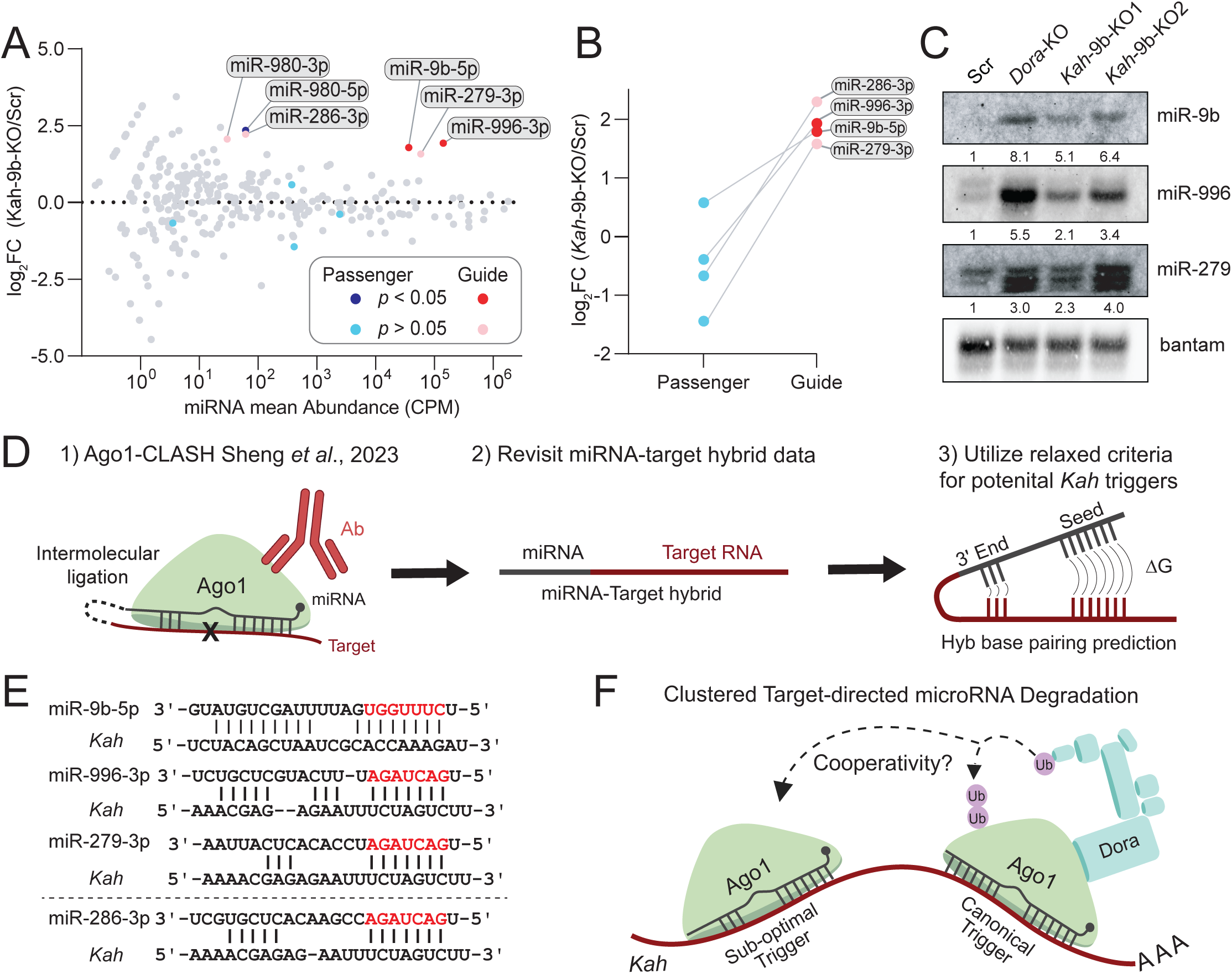
Ago1-CLASH suggests a cluster of TDMD triggers in *Kah*. **(A**) The impact of *Kah*-9b trigger knockout on miRNA abundance as determined by small RNA-seq reported in Sheng *et al.*, 2023^30^. miRNA abundance (x-axis) represents the mean counts between Scr and *Kah*-9b KO libraries, the y-axis represents the change in abundance following trigger KO in log_2_ fold-change (log_2_FC). Each dot represents an individual miRNA. Upregulated guide strands are marked in red (significant) and pink (not significant), with the passenger strands of these miRNAs being highlighted in navy blue (significant) and cyan (not significant). P-values were calculated using DEseq2. An FDR-adjusted p-value <0.05 was used as the significance threshold for this analysis (n=2 biological replicates). (**B**) A comparison of the log_2_FC between miRNA guide and passenger strands following loss of *Kah*-9b trigger. (**C**) Northern blot validating miRNA change in abundance following *Kah*-9b trigger KO. Lane labels correspond to S2 cell lines: WT (wild type), Scr (scramble/non-target sgRNA control), *Kah* trigger KO, and Dora KO. The bantam miRNA was used as a loading control as it is not sensitive to TDMD. (**D**) Reutilization of the Ago1-CLASH dataset reported in Sheng *et al*., 2023. All miRNA-target hybrids for the miR-279 family were considered when screening for potential non-canonical *Kah* triggers. (**E**) A summary of the *Kah* triggers for miR-9b, 279, and 996 found using Ago1-CLASH. miR-286 did not form hybrids with *Kah* and was therefore separated from the others by the dashed line. Red letters indicated the miRNA seed region. CLASH-identified hybrids used the Hyb base pairing pipeline to predict the most stable miRNA-trigger base pairing conformation, whereas miR-286:*Kah* base pairs were predicted using RNAcofold. (**F**) A representative model of a clustered TDMD cooperativity model, where a canonical trigger such as *Kah*-9b may nucleate the transcript for TDMD of coupled sub-optimal triggers, such as the *Kah*-279 trigger. Lavender circles represent ubiquitin and unlabeled boxes in the Dora complex represent currently unknown Culin-RING E3 ubiquitin ligase components.

**Figure S1.**
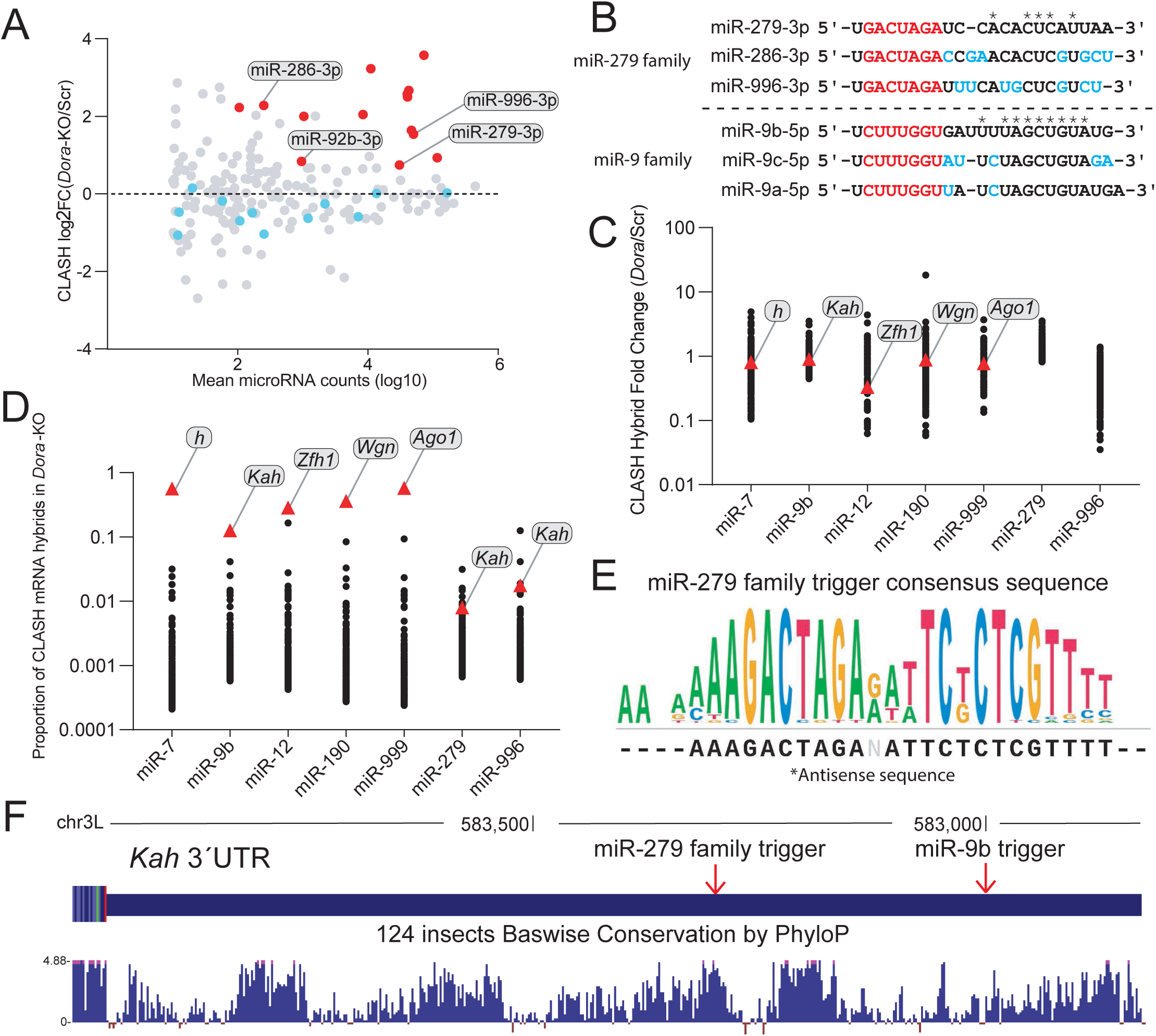
Identification of the *Kah*-279 trigger. (**A**) A comparison of the log_2_FC between guide and passenger strands of *Dora*-sensitive miRNAs found in Ago1-CLASH data from Sheng *et al.*, 2023^30^. n=2 biological replicates. (**B**) Sequence alignment of the *Drosophila* miR-279 and miR-9 families. * represent consensus nucleotides, with the seed region highlighted in red and variable nucleotides highlighted in cyan. (**C**) The fold change of individual miRNA hybrids in different miRNA cohorts in Dora-KO compared to Scr S2 cells. The most abundant hybrid for validated triggers per miRNA cohort is highlighted with red triangles and labels. Shown are the top abundant hybrids (up to 256, if able) per cohort. (**D**) The overall proportion miRNA-mRNA hybrids occupy in different miRNA cohorts in *Dora*-KO cells. Validated triggers per miRNA cohort are highlighted with red triangles and labels. Shown are the top abundant hybrids (up to 256, if able) per cohort. (**E**) The sequence conservat ion/consensus of 38 insect species. Sequence references were derived from the UCSC Genome Browser. (**F**) A PhyloP sequence conservation map (124 insects) of the *Kah* 3′ UTR derived from the UCSC Genome Browser.

We previously utilized the Ago1-CLASH method (miRNAs primarily bind Ago1 in *Drosophila*) in both *Dora*-KO cells and a Scramble (Scr) sgRNA control^30^. Ideally, CLASH ought to preferentially isolate Dora-sensitive miRNAs with their cognate triggers under *Dora*-KO conditions. To filter these miRNA-target hybrid data, we looked solely at hybrids from miRNAs that were stabilized following loss of Dora, and employed three screening criteria: 1) Extensive 3ʹ complementarity (>7 bp) separated from the seed base-pairing by at least a single mismatch (>0 nt central bulge), 2) enrichment of the hybrid following loss of Dora (>4 fold increase), 3) overall abundance of the hybrid. The miR-279 family members have divergent 3′ end sequences, meaning that a transcript that could extensively base pair with one member would be unlikely to extensively base pair with another (Figure S1B). Given that we would need to remove the extensive base-pairing criteria to screen for non-canonical *Kah* triggers, we took a closer look at how hybrid enrichment and abundance could aid our efforts in identifying them.

We initially determined hybrid enrichment based on an increase in hybrid abundance in the *Dora*-KO as compared to the Scr control^30^. However, this criterion may have potential issues as all Dora-sensitive miRNA hybrids, regardless of their contribution to TDMD, ought to increase due to the increase in miRNA abundance upon loss of TDMD. We therefore then simply looked at all the hybrids from Dora-sensitive miRNAs with validated triggers in S2 cells (miR-7, 9b, 12, 190, and miR-999) and considered the proportion each hybrid occupies in either Scr or *Dora*-KO to account for the dramatic change in miRNA abundance. To our surprise, validated triggers were not enriched when viewed in this fashion, meaning that loss of Dora merely increases the absolute number of Dora-sensitive miRNA hybrids, but did not specifically enrich bona fide targets/triggers (Figure S1C, Table S1). Thus, we next considered hybrid abundance as a readout for potential triggers. Strikingly, we found that each of our previously validated triggers, *h*:miR-7, *Kah*:miR-9b, *Zfh1*:miR-12, *Wgn*:miR190, and *Ago1*:miR-999, occupied the largest proportion of their respective mRNA hybrids even without considering base-pairing (e.g. seed-match, extensive complementarity) (Figure S1D, Table S2).

With these qualities in mind, we looked for any abundant miR-279 family hybrids with *Kah* (Figure 1D, S1D). To our surprise, we found *Kah* hybrids for both miR-279 and 996 at a highly conserved site ∼300 nt upstream of the miR-9b trigger (Figure 1E, S1E, S1F). This site did not have the extensive 3′ complementarity currently believed to be required for TDMD^26,31^; instead, the miR-996/286 3′ ends were only predicted to contain five base pairs, and miR-279 only three (Figure 1E). We did not identify any *Kah* hybrids with miR-286, again presumably due to the low expression level of miR-286 in these cells. All known transcripts that induce TDMD only contain a single site sufficient to induce decay of the associated miRNA; this secondary site against the miR-279 family would make *Kah* an endogenous example of multiple triggers clustered within its transcript, a so-called “trigger cluster”. Since this site would contain non-canonical TDMD base pairing, we considered a model where this miR-279 family trigger may be “suboptimal”, and only effective with the coupled “canonical” downstream miR-9b trigger, in a mechanism mirroring the clustered biogenesis of the suboptimal miR-451a coupled with canonical miR-144 primary transcript^43,44^ (Figure 1F). If so, then the *Kah*:miR-279 family interaction may have been disrupted by our Cas9-mediated deletion of the miR-9b trigger. Thus, we set out to interrogate the efficacy and molecular mechanism of the *Kah* trigger cluster.

### The *Kah* transcript regulates the abundance of distinct miRNA families

The most direct means for interrogating the efficacy of a potential trigger is to delete the endogenous region via CRISPR-Cas9. To do this, we generated new sgRNAs targeting either side of the putative *Kah* miR-279 family trigger to induce nucleolytic cleavage of its genomic locus (Figure 2A, S2A). Interestingly, these new KO populations (*Kah*-279-KO1/2) again showed upregulated miR-279 family and miR-9b in tandem when observed via northern blot, comparable to levels observed in a *Dora*-KO and our original *Kah*-9b-KO2 population (Figure 2B). To ensure that the observed miRNA stabilization was indeed due to loss of miRNA degradation, we employed the ‘Accurate quantification by sequencing’ (AQ-seq) method developed by Narry Kim’s group to generate minimally biased small RNA-seq libraries containing spike-in standards for count normalization^38^ (Figure 2C). In total, we found five miRNA guide strands (miR-279, 286, 996, 9b, and 92b) that were significantly upregulated (*p* adj. <.001) compared to their co-transcribed passenger strands upon loss of the miR-279 family trigger (Figure 2C, 2D). We did observe significant upregulation of the miR-980 passenger strand, though its guide was also upregulated to a similar degree, a quality likely indicative of increased biogenesis (Figure 2C).

**Figure 2.**
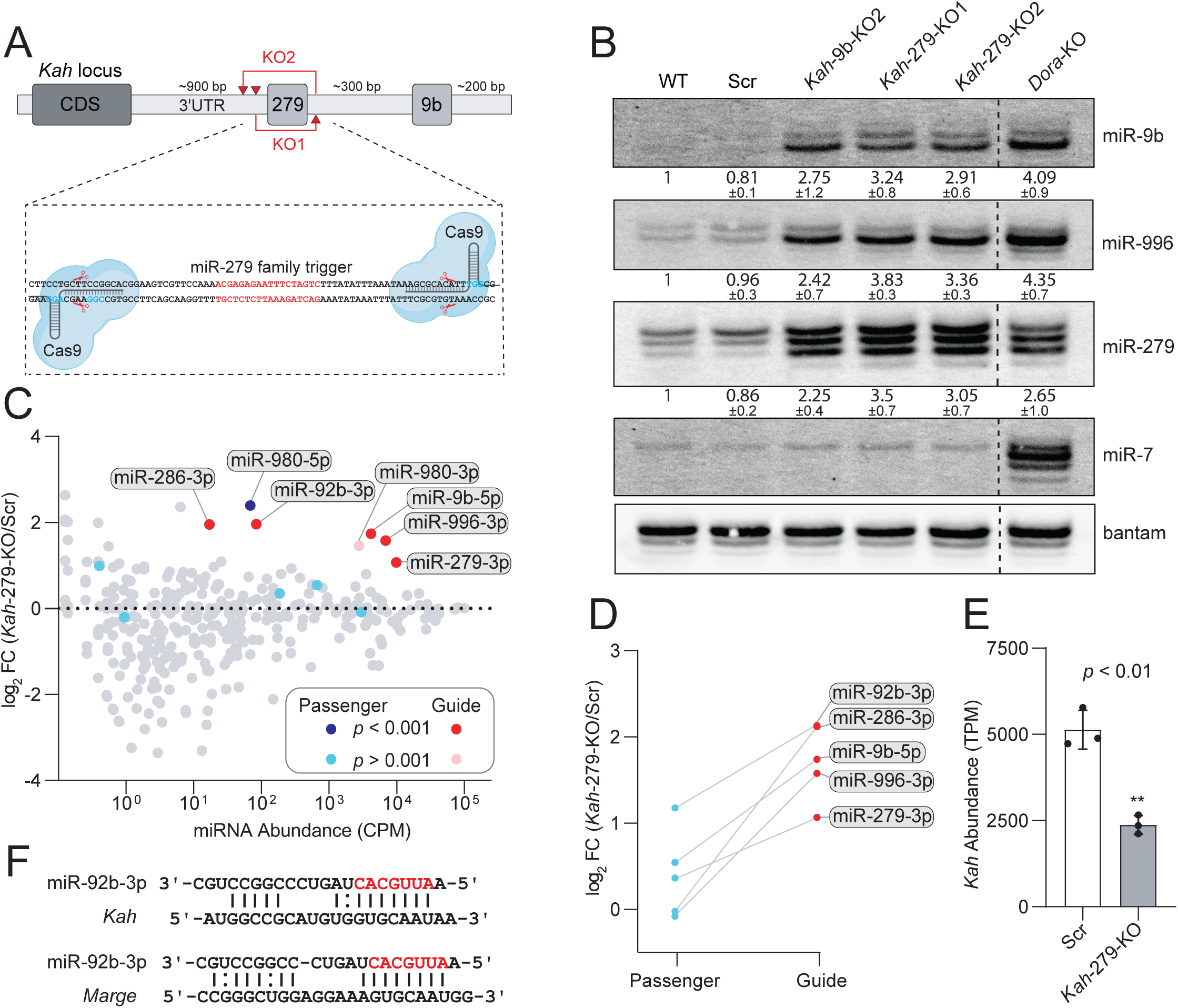
The *Kah* transcript regulates the abundance of distinct miRNA families. (**A**) A schematic of CRISPR-Cas9 targeted deletion of the *Kah*-279 trigger within the *Kah* genomic locus. Red triangles represent predicted sites of Cas9-mediated cleavage (top). The *Kah*-279 trigger sequence is highlighted in red (bottom) with the PAM sequences adjacent to each sgRNA target site highlighted in cyan. (**B**) Northern blot validating miRNA change in abundance following *Kah*-279 trigger KO. Relative miRNA levels are shown as mean ± standard deviation (SD) (n=3 biological replicates). miR-7 serves as a positive control for Dora sensitivity. ((**C**) AQ-seq detection of miRNA changes in abundance following *Kah*-279 trigger KO. miRNA abundance (x-axis) represents the mean miRNA counts per million (CPM) in Scr libraries. Highlighted are miRNAs upregulated following *Kah*-279 KO, an FDR-adjusted p-value <0.001 was used as the significance threshold for this analysis (n=2 biological replicates). (**D**) A comparison of the log_2_FC between miRNA guide and passenger strands following loss of *Kah*-279 trigger. (**E**) The influence of *Kah*-279 trigger KO on *Kah* transcript abundance as determined by RNA-seq in transcripts per million (TPM). P-values were calculated using DEseq2. ** represents a p-value < 0.01. (**F**) A potential miR-92b trigger within *Kah* identified via Ago1-CLASH, compared to the previously reported *Marge*:miR-92b interaction^21^.

**Figure S2.**
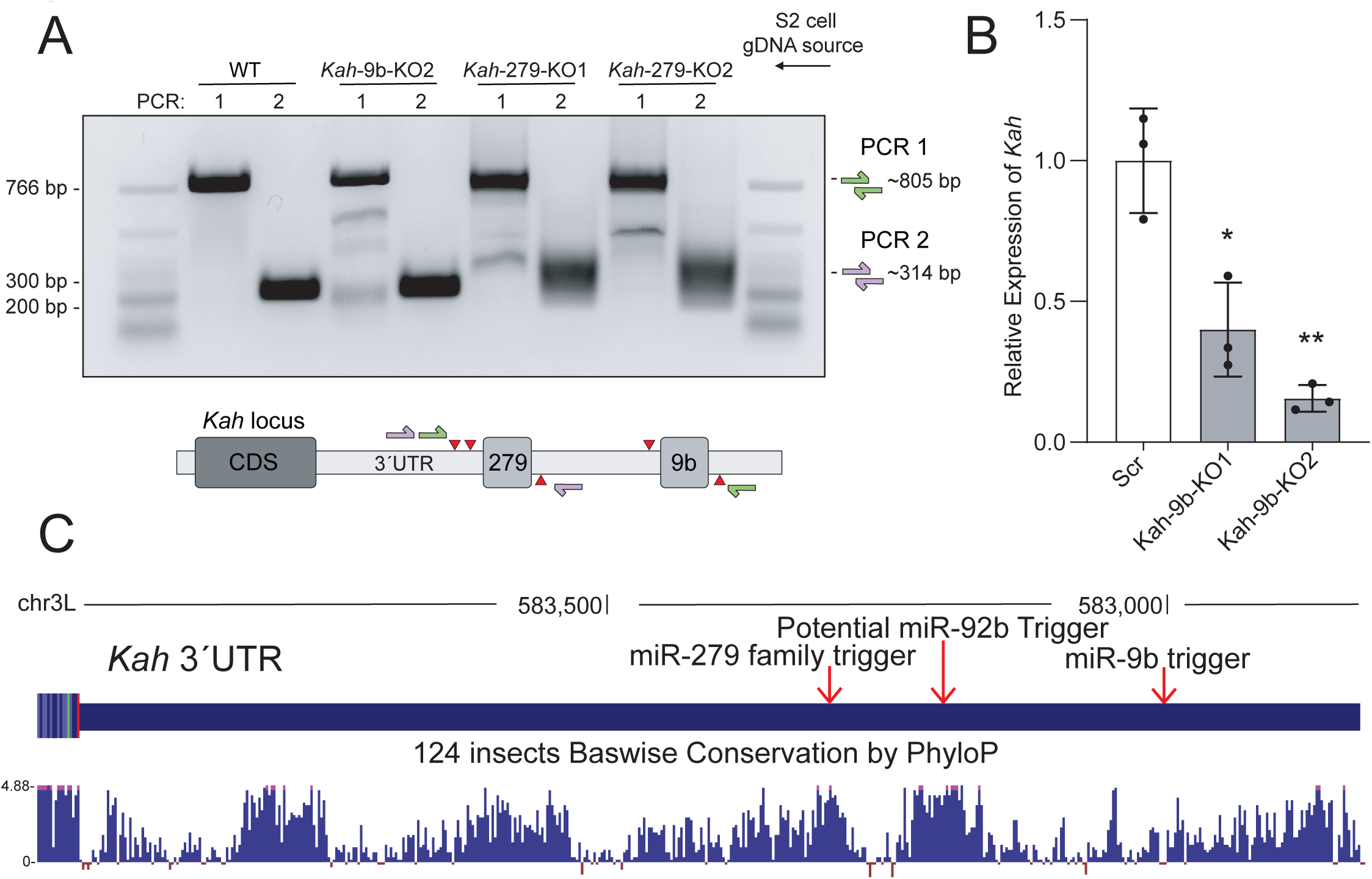
*Kah* triggers knockout validation and potential *Kah*-92b trigger. (**A**) A comparison of PCR amplicons generated from S2 gDNA. The schematic below details which primer sets were used to generate each amplicon. Destabilized loci appear less abundant, smears, and/or truncated bands. (**B**) RT-qPCR of the *Kah* transcript in *Kah*-9b KO lines. Paired t-tests, p-value *<0.05, **<0.01 n=3 biological replicates. (**C**) A PhyloP sequence conservation map (124 insects) of the *Kah* 3′ UTR derived from the UCSC Genome Browser.

While the upregulation of the miR-279 family was expected due to trigger KO, the upregulation of miR-9b and miR-92b was puzzling for a couple of reasons: 1) Under a cooperativity model, the *Kah*-279 trigger KO should only upregulate the miR-279 family, suggesting our model is likely wrong. 2) miR-9b and miR-92b are from distinct miRNA families, the miR-9 family and the miR-310 family, respectively. Several of the miR-310 family members (miR-310, 311, and 313) were reported to be sensitive to loss of Dora *in vivo* and directed for degradation by the lncRNA *Marge*^21^. However, this trigger was not observed to induce degradation of the remaining miR-310 family (miR-312, miR-92a, and miR-92b) despite them also being Dora-sensitive, suggesting that at least one other trigger exists inducing decay of the miR-310 family in addition to *Marge*^21^. An alternative hypothesis to trigger cluster cooperativity would be that CRISPR-Cas9 mediated deletions aberrantly induced *Kah* knockdown, decreasing its abundance and therefore reducing TDMD of any triggers localized within the transcript. To test this, we performed RNA-seq to monitor the change in *Kah* abundance in our *Kah*-279 KOs compared to the Scr control. Curiously, our *Kah* trigger KOs significantly reduced the abundance of the *Kah* transcript, on average having a >50% reduction, and we observed a similar trend with our original *Kah*-9b trigger KOs (Figure 2E, S2B). With this in mind, we considered whether the observed upregulation of miR-92b may indeed be due to a loss of TDMD catalyzed via another trigger within the *Kah* transcript. When reconsidering the miR-92b hybrids within our Ago1-CLASH dataset, we found a highly conserved site within *Kah* for miR-92b with only five 3′ end base pairs, a pattern mirroring what we see with miR-286/996 on the miR-279 family trigger (Figure 1E, 2F, S2C). Puzzlingly, miR-92b is predicted to base pair with *Kah* less than it is with *Marge*, yet *Marge* does not induce miR-92b degradation, suggesting that more extensive base pairing may not necessarily be more efficacious for TDMD depending on the miRNA sequence (Figure 2F). Though, due to the low expression level of miR-92b in S2 cells, it is unlikely to be an ideal model for further study of this potential trigger.

### The *Kah* trigger cluster differentially influences miRNA tailing, trimming, and function

A common feature observed of many miRNAs sensitive to TDMD is alteration to tailing (non-templated nucleotide addition) and trimming (shortening of the miRNA) on the miRNA 3′ end^10–12,16–18,24^. Tailing and trimming of these miRNAs are usually more pronounced upon loss of ZSWIM8 (or its ortholog)^16,17,24^. The major hypothesis associated with these observations is that triggers ought to extensively base pair with the interacting miRNA, potentially outcompeting the AGO PAZ domain binding of the miRNA 3′ end, leaving the end solvent exposed for non-templated nucleotide addition by TENTs or trimming by non-specific ribonucleases ^10–13,18,19,32,33^. Given the less extensive 3′ base pairing for the miR-279 family with *Kah* (Figure 1E), we sought to take a detailed look at how loss of this trigger may affect the accumulation of miRNA isoforms (isomiRs) for miR-279, 996, and 9b. To do this, we reanalyzed our AQ-seq data and categorized miRNAs based on the number of nucleotides and sequence. In this analysis, miRNAs mapping to a certain gene ought to have 100% sequence identity with the genome from nucleotides 2-18, allowing for mixed sequences at position 1 and 19-26 (Figure S3A). These criteria will capture the bulk of miRNA sequences, as it accounts for the normal size range for miRNAs, non-templated additions and trimming, as well as potential alterations in processing of the miRNA precursors by Drosha/Dicer.

**Figure S3.**
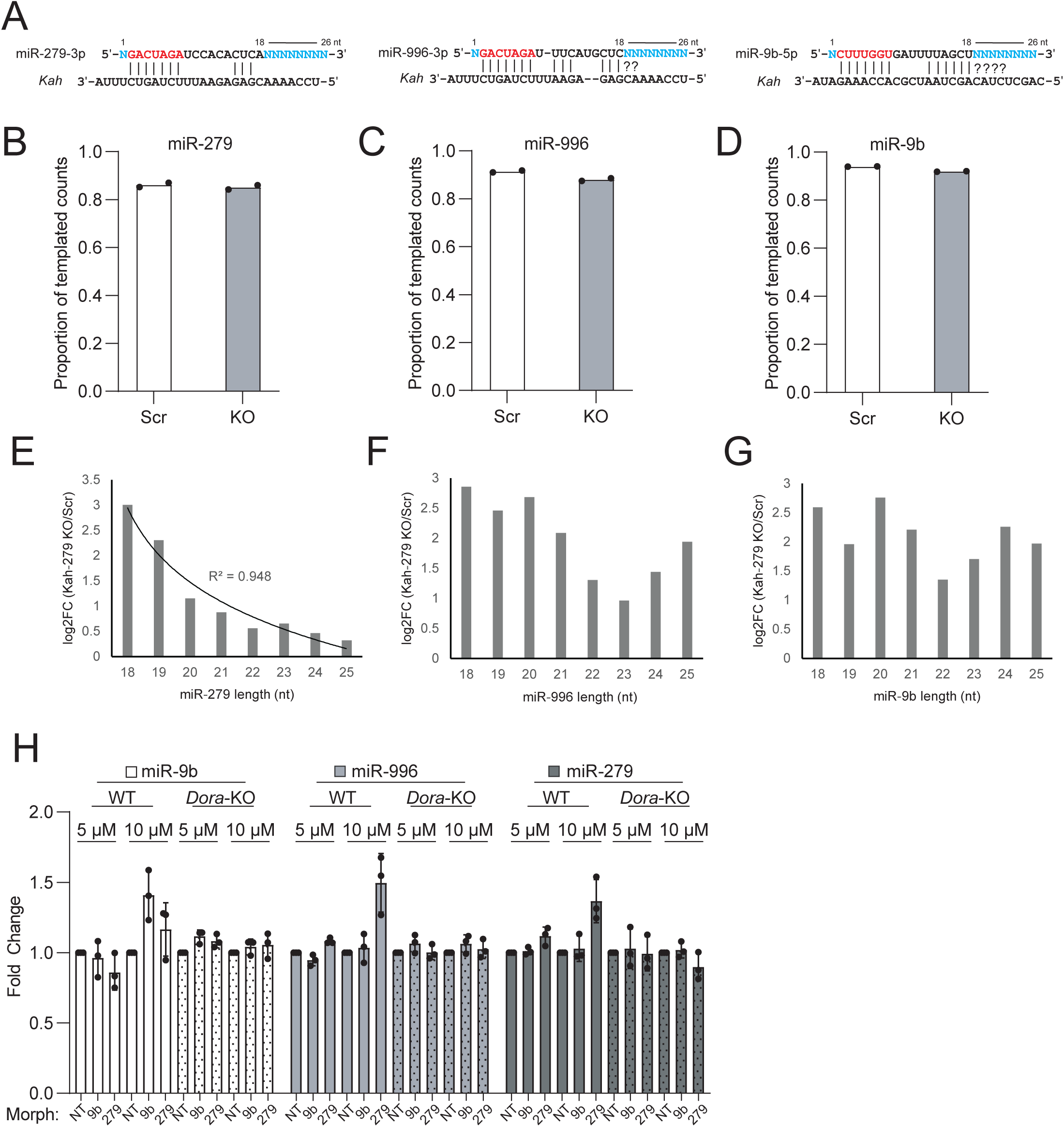
IsomiR classification and miRNA change in abundance. (**A**) A summary of our bioinformatic criteria used to classify isomiRs. miRNA seeds are marked in red, with variable nucleotide positions highlighted in cyan. Proportion of miRNA counts derived from templated sequences for (**B**) miR-279, (**C**) miR-996, or (**D**) miR-9b. n=2 biological replicates. The log_2_FC of specific miRNA isomiR lengths (*Kah*-279/Scr) for (**E**) miR-279, (**F**) miR-996, or (**G**) miR-9b. n=2 biological replicates. (**H**) Northern blot quantification of miRNA changes in abundance observed after morpholino treatment. Error bars indicate ± SD, n=3 biological replicates.

We observed a clear accumulation of shorter (trimmed) isomiRs of the miR-279 family, with little to no increase in tailing upon loss of its trigger (Figure 3A, 3B). This observation mirrors what we see for both miR-279 and miR-996 by northern, with these shorter isoforms becoming the bulk of miR-279 signal (Figure 2B). In contrast, miR-9b displayed a mix of accumulating trimmed and tailed isomiRs (Figure 3C). We next sought to define what proportion of our miRNA counts were derived from non-templated sequences. When only considering templated counts, we observed a consistent but minimal reduction in proportion of templated isomiRs, with these sequences being reduced by ∼1-3% in *Kah*-279 KOs (Figures S3B, S3C, S3D). We conclude that while loss of *Kah*-mediated TDMD alters the average length of all three miRNAs, this alteration is unlikely to be mostly mediated by tailing as a similar proportion of miRNA counts match the genomic sequence with or without the trigger. Alternatively, these observations may imply that longer-lived miRNAs may accumulate shorter/trimmed isomiRs, or even that certain templated isomiRs may be directed for turnover more easily. In line with this, we noted that shorter miR-279 isomiRs were upregulated to a much larger degree than longer isomiRs upon trigger KO, in a trend that was highly correlated (Figure S3E). Taken together, these data may suggest that shorter miR-279 isomiRs are preferential substrates for *Kah*-mediated TDMD. Even so, we did not observe a similar trend for miR-996 or miR-9b, though isomiRs shorter than 22 nt were modestly directed for turnover to a higher degree (Figures S3F and S3G).

**Figure 3.**
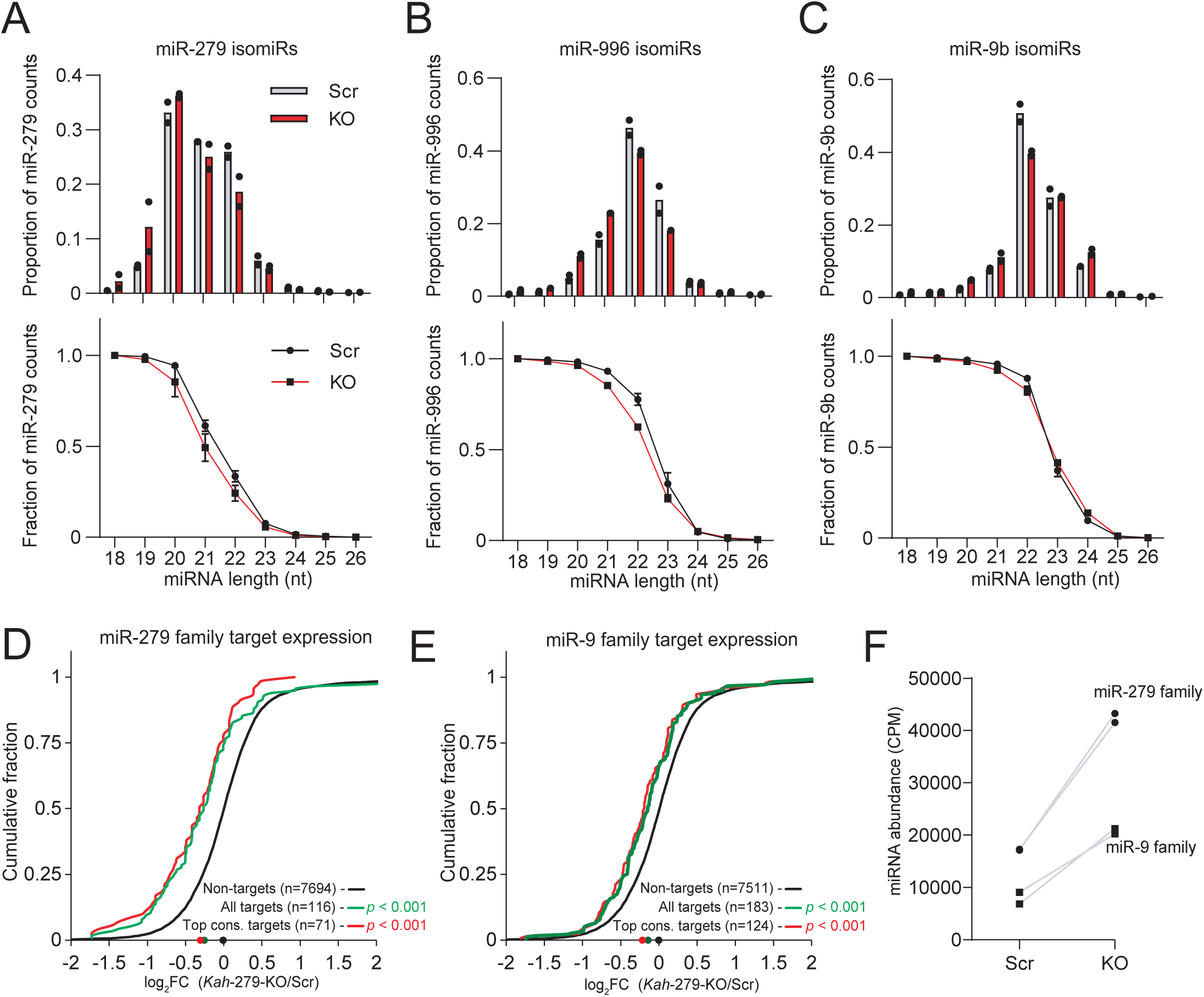
The *Kah* trigger cluster differentially influences miRNA tailing, trimming, and function. The relative proportion (top) or fraction (bottom) of isomiRs separated by length from 18-26 nts. Individual replicate values are marked with black dots (top) or an error bar (bottom). Grey and red bars represent the mean abundance in Scr and *Kah*-279 KO, respectively for (**A**) miR-279, (**B**) miR-996, or (**C**) miR-9b. The increased repression of the predicted targets of the (**D**) miR-279 family and (**E**) miR-9 family following loss of the *Kah*-279 trigger. Plotted is the cumulative change (log_2_FC) in TargetScan predicted mRNA target abundance following *Kah*-279 trigger KO compared to Scr control. The log_2_FC of individual transcripts between conditions was calculated using DEseq2. Targets were classified into all conserved targets, top conserved targets, or all other targets (non-targets) with the number of transcripts considered for each cohort listed within the plot. Dots at the bottom of the graphs represent the median expression level of each target cohort. P-values were calculated using the Mann-Whitney U test, n=3 biological replicates. (**F**) The change in miRNA family (miR-279 or miR-9) abundance following *Kah*-279 trigger KO. Individual replicates are listed as black dots (miR-279 family) or black squares (miR-9 family), n=2 biological replicates.

In any case, loss of *Kah*-mediated TDMD for the miR-279 family and miR-9b dramatically increases the overall abundance of these miRNAs, and therefore ought to increase their ability to induce repression of target mRNAs. To address this, we categorized our RNA-seq data based on TargetScan predicted mRNA targets of the miR-279 family and the miR-9 family^45^. The predicted targets of both groups were significantly repressed compared to the non-target controls, in line with the idea that the function of TDMD lies in its ability to limit miRNA-mediated silencing (Figures 3D and 3E). We did note that the overall repression of the miR-279 family targets was greater than the miR-9 family targets, a quality that becomes more understandable when considering that the absolute increase in miRNA molecules was greater for the miR-279 family (miR-279/286/996) compared to the miR-9 family (miR-9a/b/c) (Figure 3F). These results highlight how *Kah* controls the endogenous levels of distinct miRNA family targets via clustered TDMD.

### The *Kah* trigger cluster specifies miRNA decay with little crosstalk

Since the aberrant knockdown of *Kah* following trigger KOs ultimately muddles our ability to unambiguously classify predicted triggers, we next sought to further address if each trigger does indeed specify miRNA decay. To do this, we first attempted to transiently inhibit miRNA-Ago1 complex association with endogenous *Kah* using morpholino oligonucleotides (or simply “morpholinos”) against either trigger (Figure 4A). Morpholinos are short non-ionic nucleic acid analogs that can associate with RNAs based on sequence identity^46^ and have been adopted in several studies to inhibit AGO binding to targets and triggers alike^11,30,47^. This assay can help to address two outstanding questions: 1) Does the miR-279 family trigger specify decay? 2) Is the miR-279 family trigger suboptimal and dependent on the downstream miR-9b trigger? When we incubated S2 cells with 10 µM of anti-*Kah*-279 trigger morpholinos, we observed a clear stabilization of miR-279/996 in WT cells but not *Dora*-KO (Figure 4B compare lane 6 to lane 4 and lane 12 to lane 10, S3H). Interestingly, when we attempted the same experiment with anti-*Kah*-9b trigger morpholinos, we only observed stabilization of miR-9b, with no concomitant increase in miR-279/996 (Figure 4B compare lane 5 to lane 4, S3H). These changes are also unlikely to be the result of increased miRNA biogenesis or *Kah* destabilization, as we did not observe altered *Kah* or pri-miRNA abundance (Figure S4A). These results suggest that our predicted miR-279 family trigger specifies miR-279 family decay and is sufficient to recruit Dora to Ago1 on its own.

**Figure 4.**
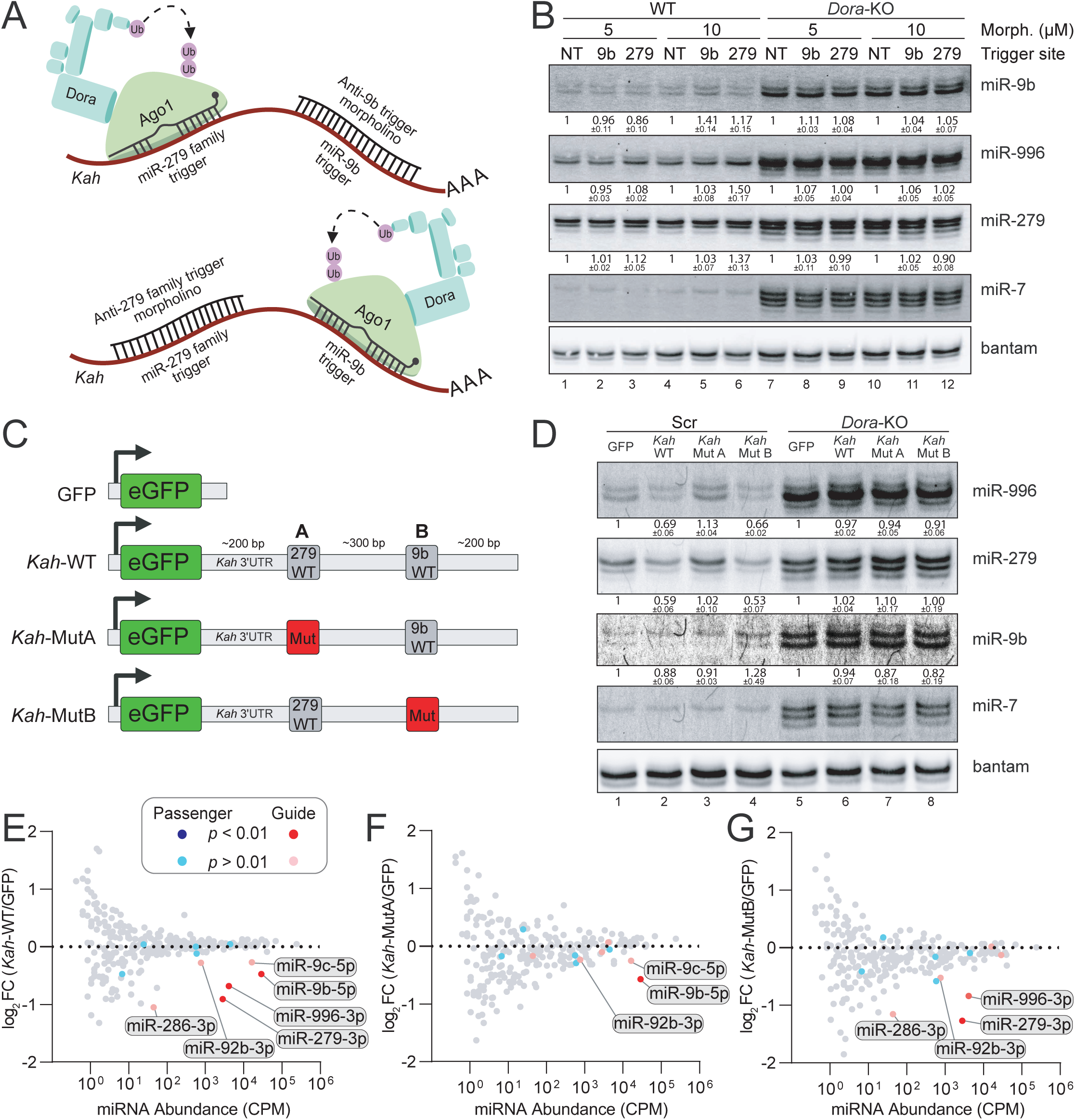
The *Kah* trigger cluster specifies miRNA decay with little crosstalk. (**A**) A schematic of anti-trigger morpholino experimental design for endogenous *Kah*. (**B**) Northern blot reporting the miRNA change in abundance following incubation with either non-target (NT), anti-9b trigger (9b), or anti-279 trigger (279) morpholinos at either 5 or 10 µM. Relative miRNA levels are shown as mean ± SD (n=3 biological replicates). (**C**) A schematic of our GFP reporter systems as described in the main text: GFP, *Kah*-WT, *Kah*-MutA, and *Kah*-MutB. (**D**) Northern blot reporting the miRNA change in abundance following introduction of reporters shown in (**C**). Relative miRNA levels are shown as mean ± SD (n=3 biological replicates). AQ-seq describing miRNA change in abundance following expression of (**E**) *Kah*-WT, (**F**) *Kah*-MutA, or (**G**) *Kah*-MutB compared to the GFP control. miRNA abundance (x-axis) represents the mean miRNA counts per million (CPM) in GFP libraries. Highlighted are miRNAs downregulated following reporter expression, an FDR-adjusted p-value <0.01 was used as the significance threshold for this analysis (n=2 biological replicates).

**Figure S4.**
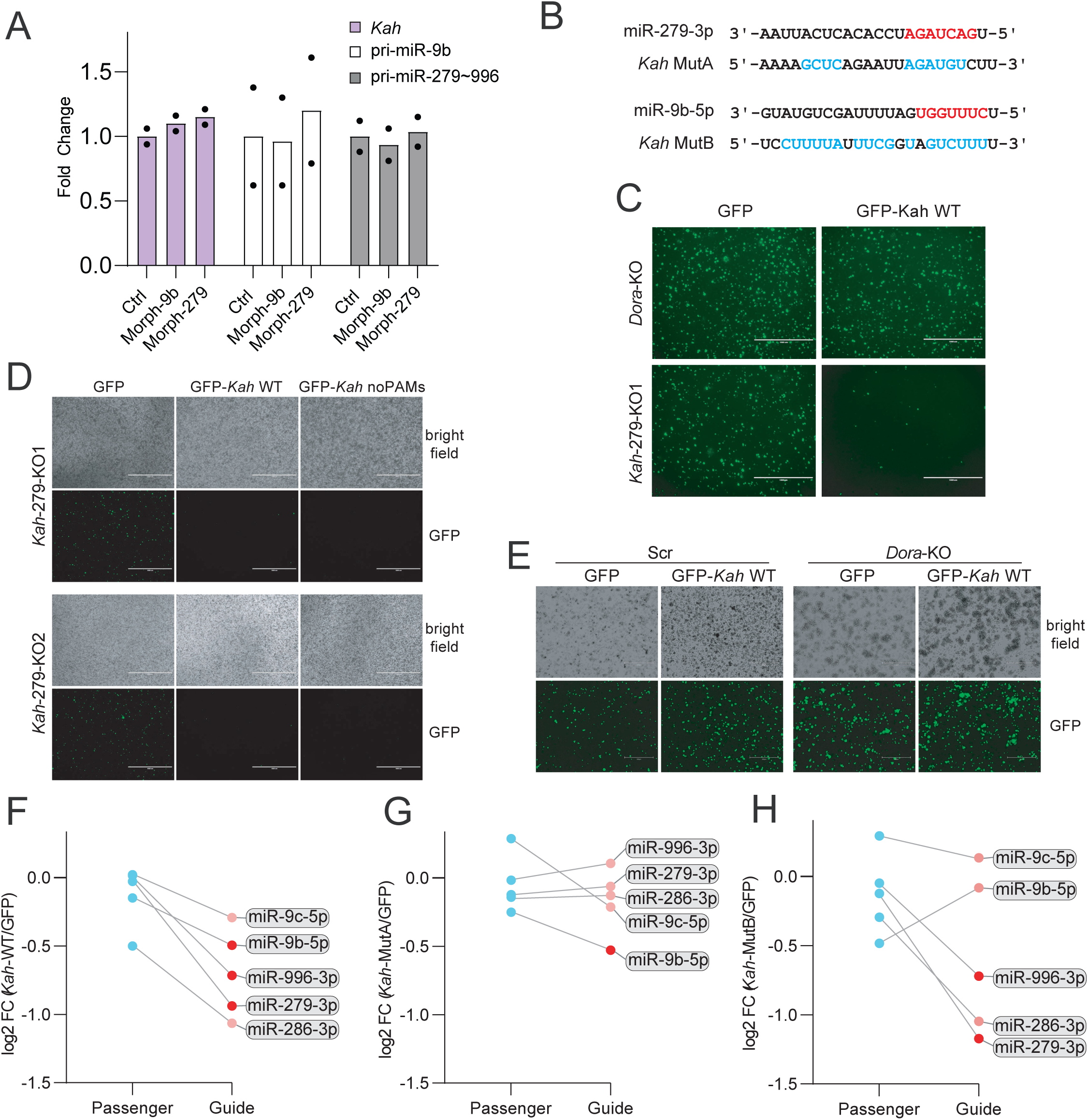
*Kah* reporter expression and miRNA strand log_2_FC. (**A**) Fold change in *Kah* or pri-miRNA transcripts in morpholino-treated cells (10µM) detected by RT-qPCR. n=2 biological replicates. (**B**) A schematic of the sequence mutations introduced in *Kah*-MutA/B reporters. Mutated nucleotides are highlighted in cyan. (**C**) GFP expression following reporter expression in *Dora*-KO or *Kah*-279-KO cells. (**D**) GFP expression in *Kah*-279 cells following transfection of *Kah* 3′ UTR constructs with or without PAM sites. (**E**) A comparison of GFP reporter expression in Scr and *Dora*-KO cells. Scale bar indicates 1000 µm. A comparison of log_2_FC observed for miRNA guide and passenger strands following expression of (**F**) *Kah*-WT, (**G**) *Kah*-MutA, or (**H**) *Kah*-MutB compared to the GFP control.

To further characterize the *Kah* trigger cluster, we next sought to induce miRNA degradation by transiently expressing a portion (∼700 nt) of the *Kah* 3′ UTR within a GFP reporter driven by a constitutively active *Drosophila* actin promoter (Figure 4C). In this experiment, we will express four constructs: GFP alone (GFP), GFP with the WT *Kah* 3′ UTR (*Kah*-WT), this same construct with mutated miR-279 family trigger (*Kah*-MutA), or an alternative with mutated miR-9b trigger (*Kah*-MutB) (Figure 4C, S4B). In this way, we can observe if each predicted trigger is sufficient to induce miRNA decay while monitoring transfection efficiency via GFP. Ideally, we would like to express these triggers in *Kah*-279 KO cells to achieve a clear and robust reduction in miR-279, 996, and 9b by northern blot. However, attempts to express constructs containing the *Kah* 3′ UTR backbone in *Kah*-279 KOs were sharply repressed compared to *Dora*-KO (Figure S4C), presumably because of the constitutive expression of CRISPR-Cas9 complex targeting *Kah*-279. We also considered generating *Kah* 3′ UTR constructs with mutated PAM sequences adjacent to the sgRNA target sites. However, these constructs were also aberrantly repressed in the *Kah*-279 KOs (Figure S4D). We instead settled on utilizing the Scr and *Dora*-KO lines for further experiments, as they offered similar robust expression of *Kah* reporters (Figure S4E). When observed by northern blot, only *Kah*-WT was able to induce miRNA decay of miR-279, 996, and 9b in tandem (Figure 4D compare lanes 1 and 2). Mutations to either trigger (*Kah*-MutA or MutB, Figure S4B) relieve repression of either miR-279/996 or miR-9b, respectively, with no observable cross-talk between the triggers (Figure 4D compare lane 1 to lanes 3 and 4). For all trigger reporters, no consistent trend of reduction was observed when expressed in the *Dora*-KO, again highlighting the Dora-dependence of these triggers (Figure 4D compare lane 5 to lanes 6, 7, and 8).

To directly quantify miRNA change in abundance induced by our reporters, we generated new AQ-seq libraries from trigger-expressing S2 cells and compared them to the GFP only control (Figures 4E-4G). We observed clear and specific repression of the miR-279 family, miR-9b/c, and miR-92b guide strands when expressing the *Kah*-WT construct in Scr S2 cells (Figure 4E). However, the only significantly changed guides (*p* adj.< .01) compared to their passenger strands were miR-279, 996, and 9b (Figure 4E, S4F). Consistent with this, it should be noted that we and the Bartel lab observed only a small increase in miR-9c following KO of the *Kah*-9b trigger, suggesting it is only minimally directed for turnover via *Kah*^21^. The overall abundance of miR-286 and miR-92b may limit reliable detection of their change responding to *Kah* reporters, given that these guides are only represented by dozens or hundreds of counts per million (CPM), respectively. Despite this, the miR-286 guide was considerably more repressed compared to its passenger strand (Figure S4F). Consistent with our northern blots, we observed no TDMD of the miR-279 family or miR-9b/c when their respective triggers were mutated in the *Kah* reporters (Figures 4F, 4G, S4G and S4H). In total, these results demonstrate that both the endogenous and ectopic expression of the *Kah* trigger cluster is sufficient to induce degradation of the miR-279 family and miR-9b.

### Structural and functional insights into the *Kah* trigger cluster

With the successful utilization of our reporter system to induce miR-279 family TDMD, we next sought to address if the minimal 3ʹ end pairing of miR-279 with *Kah* is required for turnover, or if the *Kah*-279 trigger is an example of a hypothesized “seed-sufficient” trigger^20^. At the same time, we decided to ask if the regions surrounding the *Kah* triggers may potentiate their efficacy. To answer these questions, we generated two new reporters: one with the *Kah* 3ʹ end binding sequence mutated to a seed-only pattern (*Kah*-Seed), and the other lacking the native sequences flanking each trigger (*Kah*-Short) (Figures 5A, 5B and S5A). Each of these reporters remained evenly expressed when transiently introduced into the Scr S2 cell line (Figure 5C), however both *Kah*-Seed and *Kah*-Short lost the ability to induce miR-279 family turnover, with the *Kah*-Short also losing its ability to direct miR-9b for degradation (Figure 5D). These data mirror what we have seen previously for the mammalian *BCL2L11, SSR1* and *TRIM9* triggers, in which loss of triggers’ flanking sequences in reporter constructs rendered them ineffective^29^. Together, these data suggest that the miR-279 family trigger is unlikely to be seed-sufficient, however, we cannot rule out that seemingly small mutations made to *Kah* disrupt potentially functional structural motifs, given that loss of the *Kah* trigger flanking sequences render them ineffective (Figure 5D).

**Figure 5.**
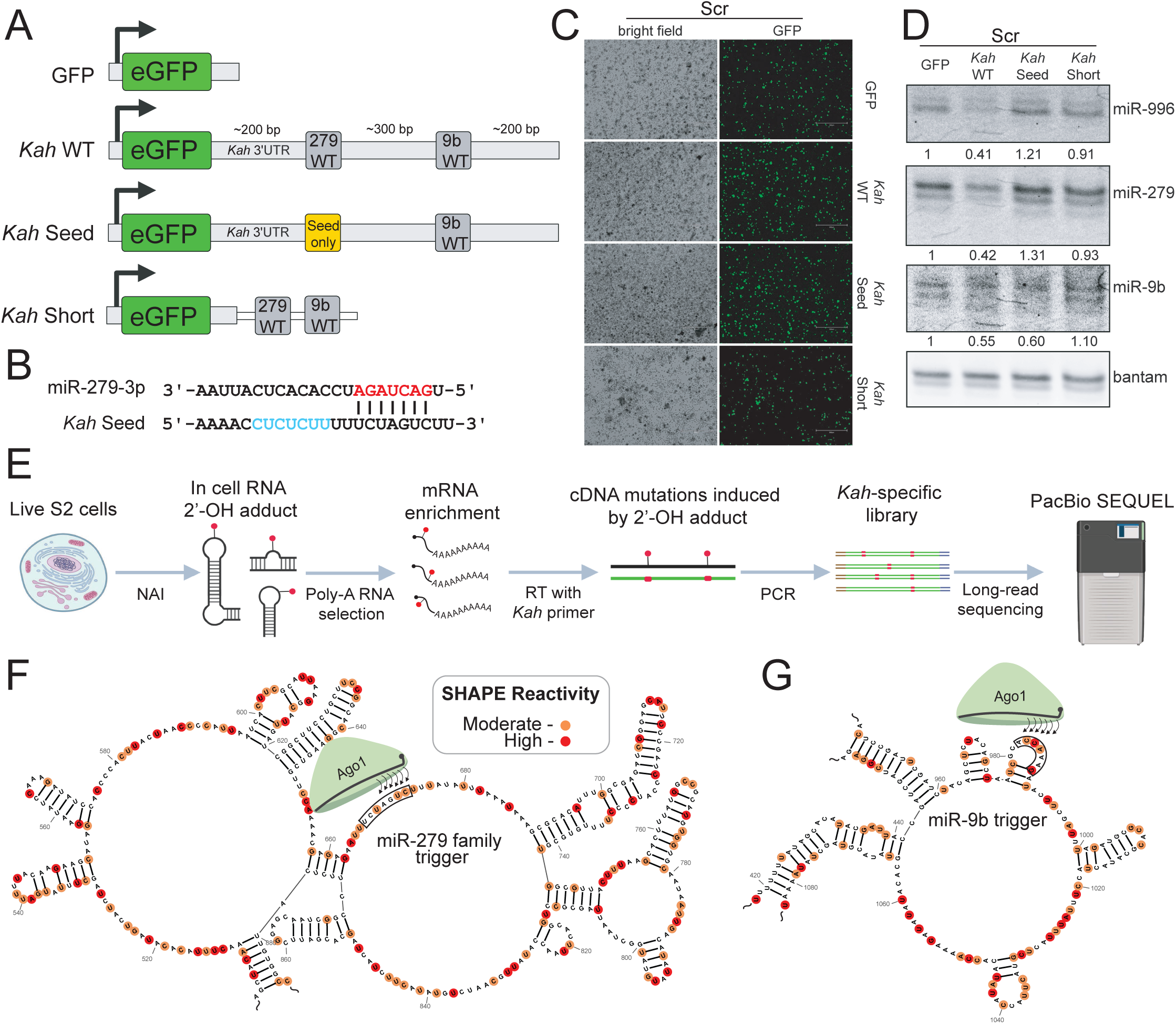
Structural and functional insights into the *Kah* trigger cluster. (**A**) A schematic of our GFP reporter system as described in the main text: GFP, *Kah*-WT, *Kah*-Seed, and *Kah*-Short. (**B**) A schematic of the predicted base pairing of the *Kah*-Seed reporter with miR-279. Red letters indicate the miRNA seed region, cyan letters indicate mutated bases. Base pairs were predicted using RNAcofold. (**C**) The transfection efficiency of the Scr cell line with each GFP reporter described in (**A**) captured with fluorescence microscopy. Scale bars indicate 300 µm. (**D**) Northern blot reporting the miRNA change in abundance following introduction of GFP reporters described in (**A**). Relative miRNA levels are shown as mean miRNA signal. (**E**) A schematic of *Kah* SHAPE library construction for long-read PacBio sequencing. (**F**) The local consensus secondary structure of the *Kah*-279, or (**G**) *Kah*-9b trigger as predicted via SHAPEmapper2. Highly reactive nucleotides are highlighted in red, with moderately reactive nucleotides highlighted in orange. Black boxes mark the seed-binding regions of either trigger.

**Figure S5.**
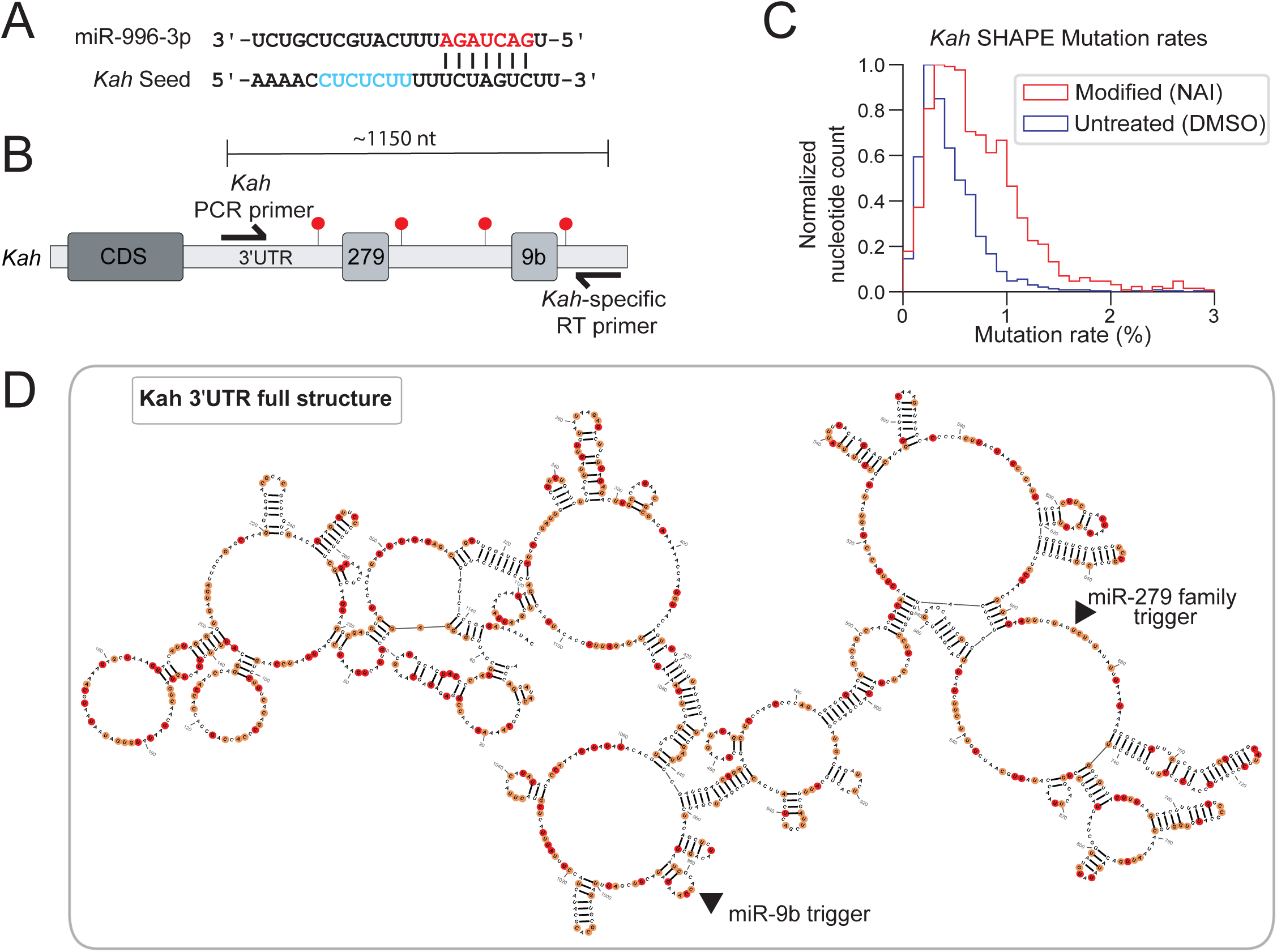
*Kah*-279 seed insufficiency and SHAPE analysis. (**A**) A schematic of the sequence mutations introduced to the *Kah*-Seed reporter. Mutated nucleotides are highlighted in cyan. (**B**) A schematic showing the region covered by long-read (∼1150 nt) *Kah* SHAPE libraries. Red points represent theoretical *Kah* bulky adducts introduced via NAI. (**C**) Mutation rates observed in DMSO and NAI-treated S2 cells calculated with SHAPEmapper2. Plotted are means derived from 2 biological replicates. (**D**) The full-length structure of *Kah* 3′ UTR derived from SHAPE-MaP libraries. Highly reactive bases are highlighted in red and moderately reactive bases are highlighted in orange.

With consistent observations of trigger flanking regions being required for TDMD, and the unique occurrence of two triggers within the same transcript, we decided to probe into the endogenous secondary structure of the *Kah* 3ʹ UTR using the Selective 2′ Hydroxyl Acylation analyzed by Primer Extension (SHAPE) method^48–50^. In SHAPE, live cells are incubated with a base modifying agent (e.g. 2-methylnicotinic acid imidazolide “NAI”) to induce chemical modification of the exposed single stranded regions of RNAs ^48–50^ (Figure 5E). These modifications can introduce mutations during reverse transcription of the cDNA (Figure 5E). cDNA libraries are then PCR amplified and subjected to high-throughput sequencing, where computational pipelines like SHAPEmapper2 can predict a consensus secondary structure based on mutational reactivity and minimum free energy^51^ (Figure 5E). To our knowledge, the only other study that has attempted SHAPE structural probing of a TDMD trigger utilized an overexpression of a *CYRANO* lncRNA fragment, given its low endogenous expression, and found that the trigger adopted a highly conserved cruciform structure ^52^. Though to what degree these results are a reflective of the endogenous *CYRANO* transcript is still unknown. Since SHAPE is primarily performed on highly abundant RNAs, we opted to enrich for *Kah* mRNAs using poly-A selection (Figure 5E). Further, since the full length of the *Kah* 3ʹ UTR exceeds 1000 nts, we took advantage of the SHAPE-competent reverse transcriptase MarathonRT popularized by Anna Pyle’s group who demonstrated its ability to reverse transcribe lower abundance RNAs as long as >2.5 kb^53^ (Figure 5E).

In the end, we settled for capturing ∼80% of the *Kah* 3ʹ UTR, in effect generating *Kah*-specific SHAPE libraries of ∼1150 nts excluding adapter sequences (Figure 5E, S5B). These long-read libraries were then sequenced on the PacBio platform. From this, we were able to perform SHAPE analysis of *Kah* using SHAPEmapper2 and observed a clear trend of increased *Kah* mutations in the NAI treated cells vs DMSO control (Figure S5C). This mutational profile was used in combination with minimum free energy calculations to generate a consensus structure of the *Kah* 3ʹ UTR (Figures S5D, S6 and S7). Using this structure, we considered the local secondary structure present at both the miR-279 family and miR-9b trigger (Figures 5F and 5G). Intriguingly, either trigger was flanked by highly structured sequences, but the seed-binding region was exposed in single-stranded portions of *Kah* (Figures 5F and 5G). In contrast, the 3ʹ end binding sequences appear locked in moderately reactive secondary structure, suggesting that these areas are primarily, but dynamically, double-stranded. These results reaffirm what has been reported for miRNA targeting dynamics previously, that miRNAs must initially associate with an accessible seed, then form a secondary duplex in the 3ʹ binding region (e.g. 3ʹ supplement), if able, before settling into its most energetically favorable conformation^1,19,54–56^. Our results suggest that the miRNA 3ʹ end binding on *Kah* may compete with local secondary structure to form a stable AGO-trigger complex, the loss of which appears to prevent miR-279 family turnover.

**Figures S6.**
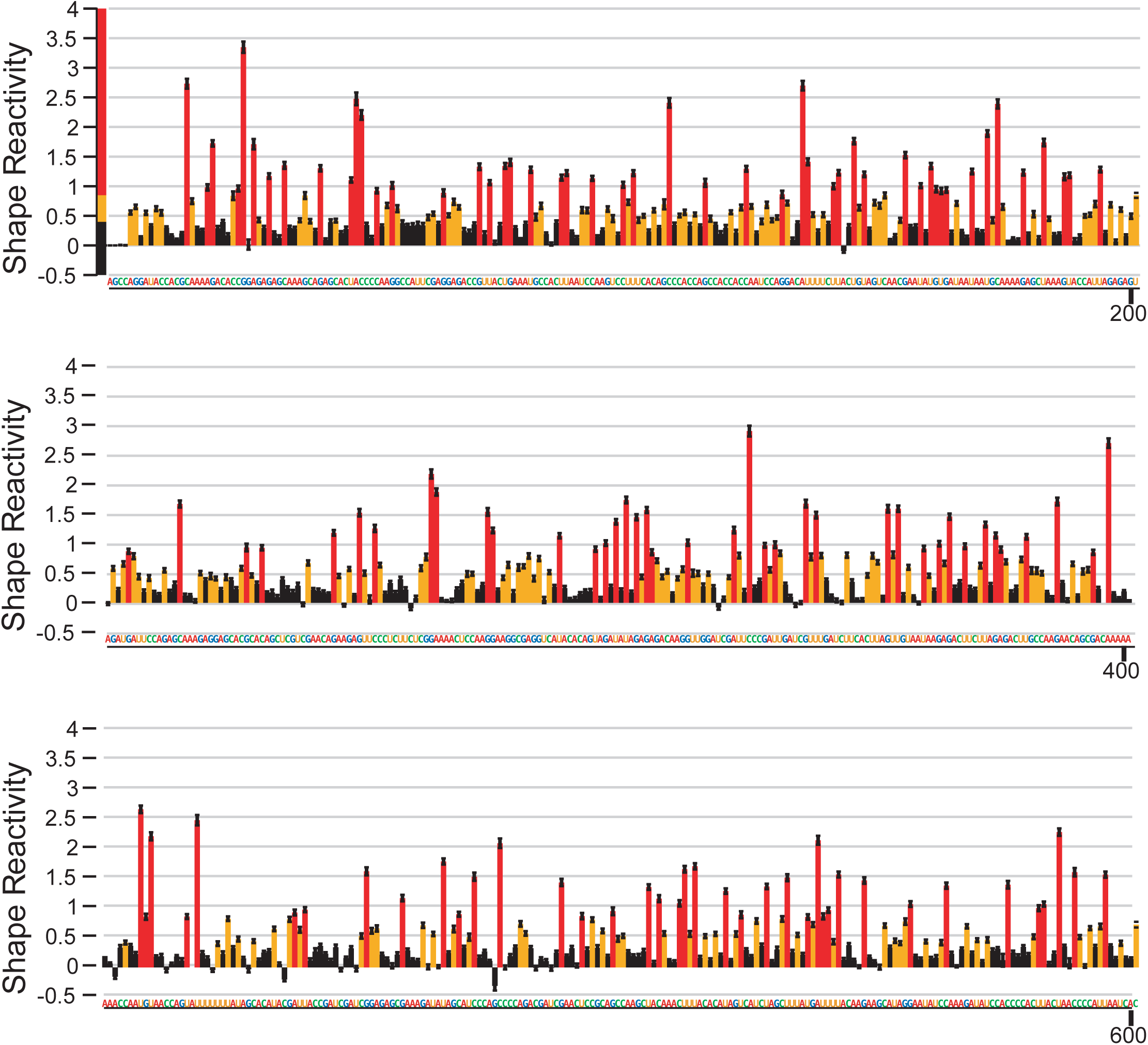
Mutational SHAPE reactivities for *Kah* (nt 1-600). Individual nucleotide reactivities for our *Kah* SHAPE libraries compared to the reference WT *Kah* transcript sequence are listed throughout. Error bars indicate SD between replicates (n=2 biological replicates). The *Kah*-279 and *Kah*-9b triggers are highlighted below their transcript locations.

**Figures S7.**
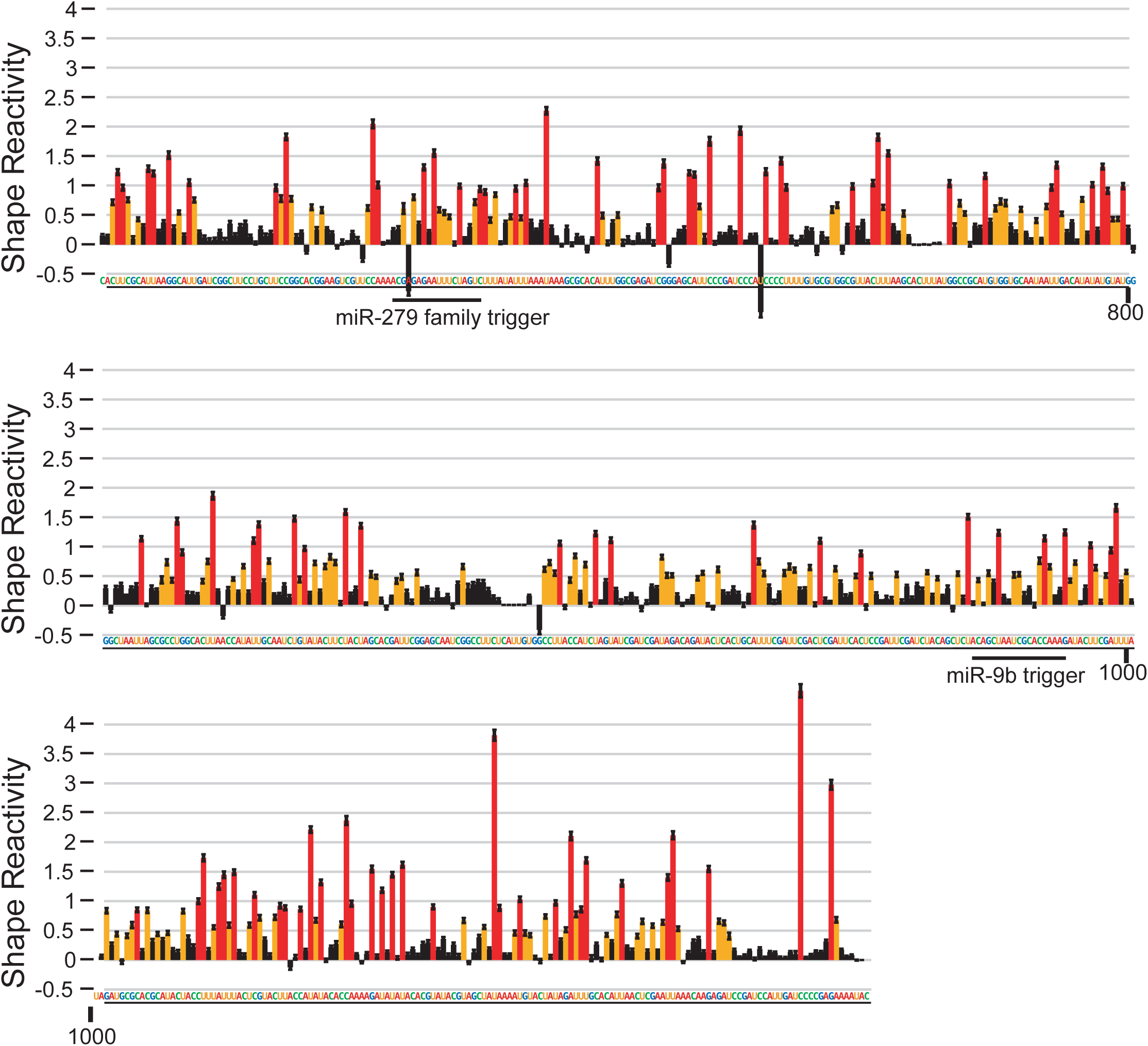
Mutational SHAPE reactivities for *Kah* (nt 601-1150). Individual nucleotide reactivities for our *Kah* SHAPE libraries compared to the reference WT *Kah* transcript sequence are listed throughout. Error bars indicate SD between replicates (n=2 biological replicates). The *Kah*-279 and *Kah*-9b triggers are highlighted below their transcript locations.

## DISCUSSION

What are the qualities that make an effective TDMD trigger? For over a decade the throughline for TDMD research has focused on a few consistent qualities across endogenous and synthetic triggers: extensive 3ʹ complementarity and seed-matching, separated from one another by a short central bulge to prevent slicing of the trigger ^26,31^. We and others are now beginning to shed light on the existence of non-canonical triggers that may require less, or no 3ʹ base pairing at all^20^. Indeed, there now appear to be three distinct classes of triggers currently identified or hypothesized: Class I triggers – require the canonical “extensive” 3′ end base pairing (≥7 bp), Class II triggers – require “minimal” 3′ end base pairing (<7 bp) such as *Kah*-279, and Class III triggers – the hypothesized seed-sufficient triggers that require no 3′ end base pairing (Figure 6A).

**Figure 6.**
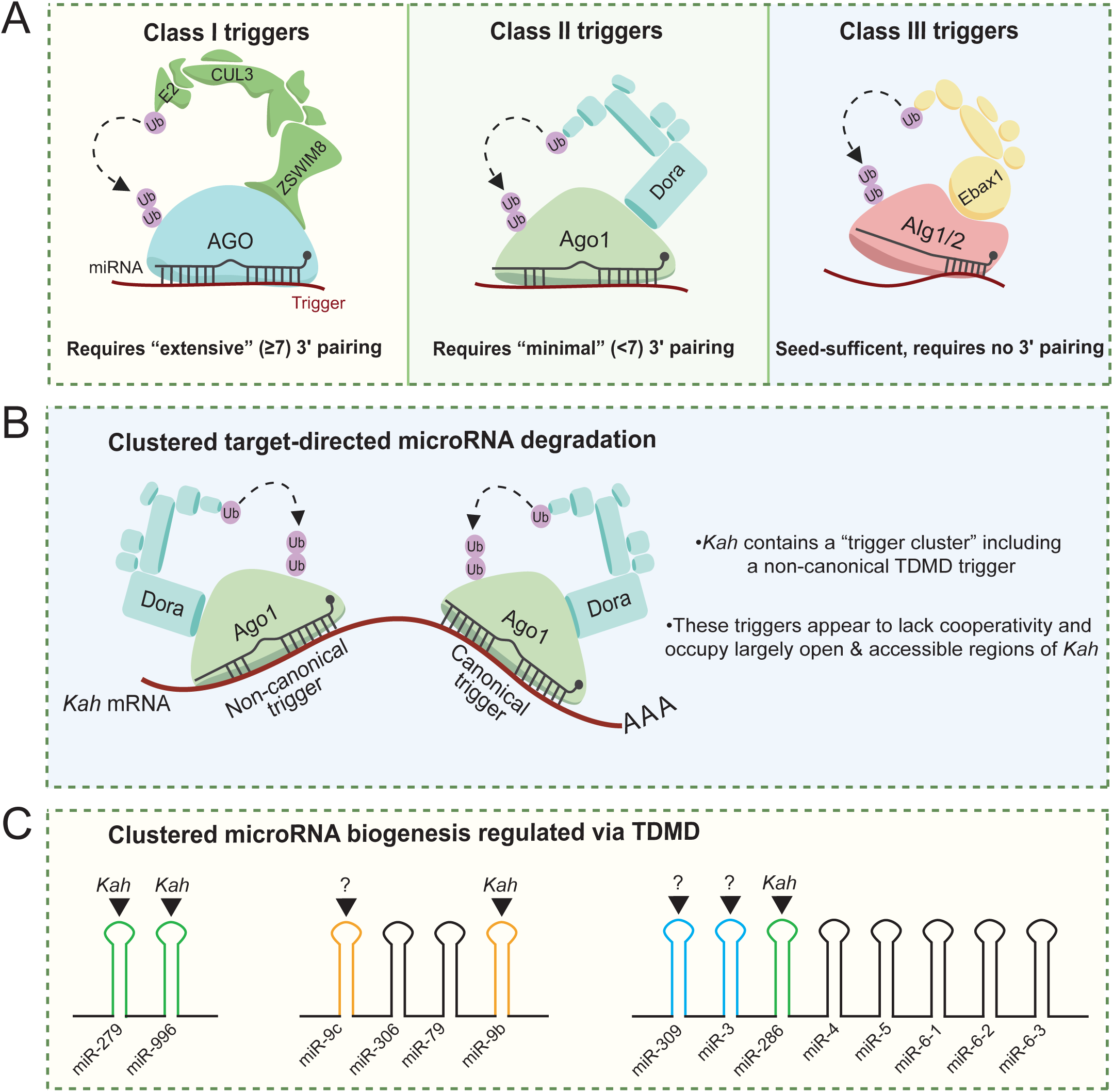
Proposed TDMD trigger classifications and clustered TDMD. (**A**) A schematic of proposed trigger classifications: Class I triggers – require “canonical” extensive 3ʹ complementarity, a representative image of the mammalian TDMD complex is shown since all known mammalian triggers currently belong to this classification. Class II triggers – require “minimal” 3ʹ complementarity, a representative image of the *Drosophila* TDMD complex is shown since *Kah*-279 fits this classification. Class III triggers – are “seed-sufficient” in that they require no 3ʹ complementarity, a representative image of the *C. elegans* TDMD complex is shown since the miR-35 family trigger is hypothesized to fit this classification. (**B**) A summary of the findings from this study: the non-canonical/Class II trigger *Kah*-279, clustered target-directed miRNA degradation, and a structural role for trigger 3ʹ complementarity. (**C**) A summary of the pri-miRNA transcripts regulated via *Kah*. Included are pri-miR-279∼996, pri-miR-9c∼9b, and pri-miR-309∼6-3. miRNAs reported to be sensitive to TDMD are indicated with black triangles above their hairpins with corresponding triggers indicated. The miR-279 family is highlighted in green, the miR-9 family in orange, and the miR-3 family in cyan.

While interesting, the existence of Class II/III triggers suggests that there must be undiscovered qualities that can effectively demarcate them as TDMD triggers and not canonical miRNA targets. The miR-279 3ʹ end interaction with guide nucleotide 15-17 (g15-17) that we observe with *Kah* is reminiscent of and partially overlaps with the canonical 3ʹ supplement (g13-16)^1^. There are several reports that point out the 3ʹ end binding sequences aid in miRNA-target recognition in part by competing with local target secondary structure^35,56–58^. Here, we used SHAPE to probe the endogenous secondary structure of the *Kah* triggers and found that while the seed-binding region was highly accessible, the 3′ end binding sites are locked in secondary structure. To what degree 3ʹ end complementarity is required for an AGO TDMD-competent conformational change, and not simply stabilizing association and/or outcompeting other miRNA targets, has still not been thoroughly investigated. Many studies reporting *in vitro* miRNA targeting assays have highlighted a reduction in AGO association with targets containing only a seed match with no 3ʹ complementarity^57,59,60^. Future research into the influence of 3ʹ end complementarity in AGO association with various triggers may help to answer these outstanding questions. Additionally, the list of validated triggers has grown considerably since the first crystal structures of an AGO-trigger complex were reported^19^. While challenging, it may be crucial to expand on these findings by generating new crystal structures spanning the validated triggers, canonical and non-canonical alike, to identify any critical structural rearrangement common between them upon trigger association.

Here, we found that the minimal miR-279 3ʹ base pairing with *Kah* was required for TDMD, but not all isoforms were directed for turnover evenly. The shortest miR-279 isoforms were degraded to a larger extent compared to longer isoforms, suggesting that miRNA length may be a contributing factor for TDMD of certain miRNAs. Interestingly, given that the miR-279 base pairs with *Kah* end at g17, and isoforms approaching g18 were turned over more quickly, there is the potential that the 3ʹ end base pairing proximity to the terminal nucleotide may contribute to TDMD. A well-known example of a highly potent canonical TDMD trigger is the *CYRANO*:miR-7 interaction, which has uniquely extensive 3ʹ complementarity (∼14 bp) for this abnormally long miRNA (∼24 nt)^12^. This interaction likely promotes the miR-7 3ʹ end exit from the AGO PAZ domain, leaving the end solvent exposed for non-templated additions via Gld2 (TENT2, TUT2), though the tailing of miR-7 during engagement with *CYRANO* is dispensable for TDMD^12^. The *Kah* trigger cluster does not appear to readily induce tailing of either the miR-279 family or miR-9b, despite miR-9b having the canonical extensive 3ʹ base pairing typical of triggers. However, removal of the *Kah* trigger did allow for the accumulation of shorter, templated, miRNA isoforms for both the miR-279 family and miR-9b. These results highlight that tailing and trimming may be exceptionally dynamic based on the miRNA, trigger RNA, 3ʹ end base pairs, and the model system used for study.

Despite recent successes in identifying endogenous triggers, there are still many left to be found. AGO-CLASH and similar methods appear to be a powerful tool for capturing trigger RNAs interacting with their cognate miRNAs. Here, we established that highly abundant miRNA-target hybrid reads in our Ago1-CLASH datasets were predictive of *bona fide* triggers, aiding in our screen for non-canonical triggers within *Kah*. These results agree well with another report in *Drosophila* suggesting that the abundance of TDMD triggers may be a key determinant of efficacy^21^. Validation of the non-canonical *Kah*-279 trigger revealed that the *Kah* transcript contains a cluster of triggers, each inducing miRNA degradation independently of the other (Figure 6B). Since *Kah*-279 is sufficient to induce decay of all the entire miR-279 family, it was more potent in its ability to influence miR-279 targets compared to miR-9 targets upon loss of *Kah*-mediated TDMD. These results highlight how clustered TDMD provides an additional layer of complexity to the tissue-specific gene regulation, in that tissues expressing *Kah* may also express the miR-279 family and miR-9b, only one of these miRNAs, or none of these. Such combinations would imply that *Kah* could simply act as an mRNA, a trigger, or a trigger cluster depending on the context. The idea of clustered TDMD also highlights how it was recently shown that the primary transcripts of miRNAs directed for TDMD in mammals preferentially organize into clusters, with TDMD acting as a tool for cells to augment the abundance of select miRNAs derived from polycistronic transcripts^24,25^. Interestingly, *Kah* induces decay of the miR-279 family in addition to miR-9b, suggesting *Kah* post-transcriptionally augments the abundance of several miRNAs derived from three distinct polycistronic transcripts (Figure 6C).

The only other, to our knowledge, reported attempt to express a cluster of triggers to degrade miRNAs predates the discovery of ZSWIM8’s role in catalyzing TDMD^13^. In contrast to our study, they used multiple (4x) canonical triggers, directly adjacent to one-another, against single miRNAs within the same transcript and found that the additional sites reduced TDMD efficacy compared to single trigger^13^. In the endogenous context, there may be a “sweet spot” in the number of triggers, and the spacing between those triggers, that can exist within the same transcript. Consistently, we found that several triggers require flanking regions for them to be effective^29^ (Figure 5). More broadly, expanding the list of endogenous triggers may aid in the development of potent synthetic triggers able to induce decay of individual miRNAs, entire miRNA families, or several miRNA families via clustered TDMD. Such tools would be key for probing the influence of specific miRNAs during development and disease alike.

While we provide a detailed molecular analysis of the *Kah* trigger cluster in S2 cells, how integral the TDMD of these miRNAs are to the *Drosophila* lifecycle will be the subject of future research. Interestingly, *Kah* (*Kahuli*) is predominantly expressed in the *Drosophila* mesoderm during development^61^, but the miR-279 and miR-9 families are often neuronally expressed with some exceptions^62–65^. There are reports of the miR-279 family being integral to the development of *Drosophila* sensory organs, with miR-279 and 996 likely functioning redundantly^63,64^. In line with this, we find that not only do miR-279 and 996 target redundantly, but they also appear to be concurrently degraded. Further *in vivo* analyses would be key in providing a spatiotemporal view of *Kah* expression, miR-279 family and miR-9b TDMD, and any phenotypic consequences therein.

## STAR*METHODS

### EXPERIMENTAL MODEL AND SUBJECT DETAILS

#### Cell lines and cell culture

*Drosophila* S2 cells were maintained in Schneider’s Insect Medium (Sigma, S9895) supplemented with 10% heat-inactivated FBS (Cytiva, SH3054103HI) and 1% penicillin-streptomycin (Gibco, 15140163) at 28°C. Cells were passaged 1:5 every 4-5 days when the culture reaches confluency.

### METHOD DETAILS

#### Ago1-CLASH prediction of non-canonical *Kah* triggers

All miRNA-RNA hybrids were extracted from previously reported Ago1-CLASH data^30^. These hybrids were summarized based on miRNA and target sequences, target RNA type (e.g. mRNA, rRNA, lncRNA), and mean abundance of this hybrid in Scramble and *Dora*-KO cell replicates (Tables S1 and S2). The scripts used to perform these analyses are available on GitHub (https://github.com/UF-Xie-Lab/TDMD-in-Drosophila). From this summary, hybrids were manually reorganized based on miRNA identity (e.g. miR-999 hybrids, miR-279 hybrids), and sorted based on mean hybrid abundance in either Scramble or *Dora*-KO cells. Hybrid enrichment in the *Dora*-KO was assessed by taking each miRNA hybrid cohort (e.g miR-279 hybrids) and determining the overall proportion each individual hybrid occupied within the cohort to account for miRNA change in abundance following *Dora*-KO. Fold changes between the Scramble and *Dora*-KO were compared to assess hybrid enrichment (Table S1). The abundance of individual hybrids in *Dora*-KO, as this data should contain the most abundant Dora-sensitive miRNA-RNA hybrids, was similarly reorganized into miRNA cohorts. The overall proportion each individual hybrid occupied within the cohort was calculated, with non-mRNA hybrids being excluded to remove spurious miRNA hybrids (e.g. miRNA-miRNA, miRNA-rRNA, or miRNA-tRNA hybrids) (Table S2).

#### Plasmid construction

*Drosophila* gene-specific knockout plasmids were generated using the pAc-sgRNA-Cas9 (Addgene, 49330) vector. Three sgRNA expression plasmids were used in pairs flanking the *Kah*-279 trigger sequence. sgRNA sequences inserted into the expression vector are as described in the supplementary table (Table S3). For GFP and GFP-*Kah* expression vectors, the pAC-sgRNA-Cas9 (Addgene, 49330) plasmid was used as a backbone wherein the Cas9 CDS was swapped for GFP with or without insertion of the *Kah* 3ʹ UTR. The *Kah* 3ʹ UTR was amplified by PCR from genomic DNA extracted from S2 cells using standard organic nucleic acid extraction. Mutants at specific *Kah* locations were introduced via overlap extension PCR. Primers for the construction of these vectors are listed in the supplementary material (Table S3).

#### Transfection and stable cell line generation

Transfection of S2 cells was performed according to the manufacturer’s protocol using either lipofectamine 3000 (Invitrogen, L3000015), or the SF Cell Line 4D X Kit (Lonza, V4XC-2024) on the 4D-Nucleofector X Unit (Lonza, AAF-1003X) platform using the default S2 cell settings. All transfections were performed using 1 μg plasmid/1×10^6^ S2 cells and allowed to grow for 72 hours following transfection before being collected. GFP signal was captured from reporter-transfected S2 cells using either an EVOS FL (Invitrogen, AMF4300) or EVOS M5000 (Invitrogen, AMF5000). Stable lines were generated by antibiotic selection using 5 μg/mL puromycin (Gibco, A1113803) for at least 3 weeks.

#### Morpholino oligonucleotide treatment

S2 cells were passaged normally prior to introduction of custom vivo-Morpholino oligos (GeneTools, Table S3) which was performed as described previously^30^. Briefly, 6×10^6^ S2 cells/well were seeded into 6-well plates and incubated with either non-target, Anti-279-trigger, or Anti-9b-trigger vivo-morpholinos at either 5 or 10 μM. The cells were then cultured for additional 48 hrs, at which point cells were collected for total RNA extraction.

#### RNA isolation and RT-qPCR

To collect total RNA, cells were pelleted by centrifugation at 300xg for 2-3 minutes, washed once with 1X phosphate buffer saline (PBS) (Gibco, 10010023) and the cell pellets were resuspended in and extracted using the TRIzol Reagent (Invitrogen, 15596018) following the manufacturer’s protocol. For RT-qPCR experiments, cDNA was generated from total RNA samples using the QuantiTect Reverse Transcription Kit (QIAGEN, 205313). qPCR experiments were performed using SsoAdvanced Universal SYBR Green Supermix (Bio-Rad, 1725275) and data was normalized to *Actin*. All primers used in this study are listed in Table S3.

#### Northern blot

Near-infrared northern blots were performed as previously described^29,30,42^. Briefly, 10-20 µg of total RNA per sample was separated on a 20% denaturing polyacrylamide 7M urea gel. RNA was transferred (semi-dry) to the Nytran N (Cytiva, 10416196) nylon membrane. Blots were then chemically crosslinked to the membrane via 1-Ethyl-3-[3-dimethylaminopropyl]carbodiimide hydrochloride (EDC) (Thermo Scientific, 22981). Crosslinking reagents were then rinsed off the membrane with water, and preincubated in ExpressHyb hybridization buffer (Takara, 636833). Blocked membranes were then incubated with IR-dye labeled antisense oligonucleotide probes or azide-labeled oligonucleotides which can be conjugated to IR dyes (Table S3) against the desired RNAs for at least 6 hours. IR signal was captured with an Amersham Typhoon (Cytiva, 29238583) and images were analyzed with ImageQuant TL (v7.0).

#### RNA-seq library preparation

Total RNA samples were depleted of genomic DNA contamination using Turbo DNase (Invitrogen, AM1907) according to the manufacturer’s protocol. RNA quality was assessed via Agilent TapeStation (Agilent, G2992AA) on RNA ScreenTapes (Agilent, 5067-5576). RNA samples with an RNA integrity number (RIN) above 9 were used for library preparation. RNA sequencing was outsourced to MedGenome for library prep using the KAPA mRNA HyperPrep kit (Roche, 08098123702) and subsequently sequenced on the Illumina NovaSeqX Plus platform.

#### AQ-seq library preparation

AQ-seq small RNA libraries were generated as described previously^38^. Briefly, 10-20 µg of total RNA per sample was mixed with 1 µL 3.33 nM synthetic spike-in small RNAs and was size selected for small RNAs on a 15% polyacrylamide urea gel. Gel purified small RNAs are ligated to a pre-adenylated 3′ adapters with RNA Ligase 2 KQ (NEB, M0373L). Ligated small RNAs were again size selected via urea-PAGE. 3′ ligated small RNAs were then ligated to the 5′ adapter with RNA Ligase 1 (NEB, M0437M). Following the final ligation, small RNAs were directly reverse transcribed using SuperScript III Reverse Transcriptase (Invitrogen, 18080085). cDNA libraries were amplified using NEBNext High-Fidelity 2X PCR Master Mix (NEB, M0541L) and library sizes/concentrations were estimated via Agilent Tapestation (Agilent, G2992AA) on DNA ScreenTapes (Agilent, 5067-5582). Libraries were sequenced either by Admera Health or the University of Florida Interdisciplinary Center for Biotechnology Research (ICBR). Adapters, primers, and spike-in sequences are listed in Table S3.

#### *In vivo* RNA SHAPE modification

*In vivo* S2 RNA modification was performed as previously described with minimal modifications^53^. Briefly, WT S2 cells were passaged normally for 2-4 days prior to collection to ensure cells were growing in log phase. Per SHAPE replicate, 3×10^7^ cells were collected and resuspended in the SHAPE modification mixture (0.4 U/μL SUPERaseIn [Invitrogen, AM2694], 200 mM NAI [Millipore Sigma, 03-310]) brought up to 3 mL total volume with 1X PBS. Control samples were collected in tandem with the NAI volume being swapped for 100% LC-MS grade DMSO (Thermo Scientific, 85190). Cells were incubated in the mixture at room temperature (∼24°C) for 10 minutes turning constantly (10 RPM), pelleted at 250xg for 2 minutes, the supernatants were removed, and pellets were resuspended in 2 mL TRIzol Reagent (Invitrogen, 15596018) per replicate. Biological replicates were performed on separate days using fresh materials. Total RNA from these cells was collected as previously described.

#### Poly-A RNA selection

To enrich *Kah*, poly-A selection was performed on SHAPE-modified (NAI) and control (DMSO) RNA samples using Dynabeads Oligo (dT)_25_ (Invitrogen, 61005) using the manufacturer’s protocol with minor modifications. 225 μg total RNA (per condition/replicate) was resuspended up to 300 μL with 10 mM Tris-HCl, pH 7.5. 300 mg of beads were used per condition/replicate. Beads were washed with 1 mL of Binding Buffer (20 mM Tris-HCl, pH 7.5; 1.0M LiCl; 2 mM EDTA), separated onto a magnet stand, and the supernatant was removed. Beads were then resuspended with Binding Buffer equal to the original volume of beads aliquoted. An equal volume of Binding Buffer was added to diluted total RNA (300 μL), and briefly mixed. Washed beads were added to total RNA and mixed by gentle agitation briefly. Bead/total RNA mixture was incubated in a pre-heated thermomixer set to 80°C for 3 minutes, then allowed to cool down slowly to 37°C (∼10-15 minutes). Tubes were then placed onto magnet stand, and poly-A depleted supernatant was discarded. Beads were washed with 600 μL of Washing Buffer B (10 mM Tris-HCl, pH 7.5; 0.15M LiCl; 1 mM EDTA) by pipetting. Tubes were placed on the magnet stand and washed again, with the supernatant discarded. Beads were then resuspended in 60 μL of 10 mM Tris-HCl, pH 7.5. The samples were then heat-denatured at 77 °C for 2.5 minutes to elute poly-A RNAs off the beads. Eluate was separated from the beads on a magnet stand, eluted poly-A RNAs were ethanol precipitated and monitored via high-sensitivity RNA ScreenTape (Agilent, 5067-5579) before continuing with SHAPE library preparation.

#### Long-read *Kah* SHAPE library construction

To generate the cDNA for long-read *Kah* SHAPE libraries, the UltraMarathon Reverse Transcriptase (uMRT) kit (RNAConnect, N/A) was used according to the manufacturer’s recommendations. In brief, a fresh uMRT master mix (50 mM Tris-HCl, pH 7.5; 200 mM KCl, 1 mM MnCl_2_, 20% Glycerol, 0.25 μM *Kah*-RT primer, 0.25 mM dNTPs, 1U/μL uMRT Enzyme, 1U/μL SUPERaseIn, 2.5 mM DTT) was mixed with the Poly-A RNA samples and allowed to reverse transcribe for 3 hrs at 42°C. cDNA was cleaned up from short primers via 1X AMPureXP (Beckman Coulter, A63881) according to the manufacturer’s protocol. cDNA was eluted and amplified using NEBNext High-Fidelity 2X PCR Master Mix (NEB, M0541L) and indexed primers until sufficient material was obtained for PacBio library construction. Library size and concentration was monitored via D5000 ScreenTape (Agilent, 5067-5588). PCR amplicons were sent to the University of Florida ICBR for PacBio SMRTbell library construction and sequenced on the PacBio Sequel IIe system.

### QUANTIFICATION AND STATISTICAL ANALYSIS

#### Northern blot quantification

Northern blot images were quantified using ImageQuant TL (v7.0). In brief, raw microRNA signal was quantified, and the background signal was subtracted from each value. Relative microRNA quantification was then normalized to the bantam loading control. Normalized values were then compared to the negative control lane. Blots with ± values represent the normalized signal standard deviation between 3 biological replicates.

#### RNA-seq analyses

The RNA-seq differential expression analyses follow a standard pipeline. After adapter trimming using Cutadapt^66^, the clean reads were aligned to the reference genome using HISAT2 to generate a mapping file^67^. Gene-level read counts are then computed using the prepDE.py3 script. Subsequently, the read counts were input into the DESeq2 software for differential expression analyses, enabling the identification of differentially expressed genes (DEGs) between conditions^68^. For miRNA family target analyses, these data were categorized based on conserved miRNA targets predicted through TargetScanFly (v7.2)^45^. Top conserved targets were separated from all conserved targets based on a criterion of <-.40 conservation context score.

#### AQ-seq analyses

The analysis of AQ-seq data began with preprocessing raw sequencing reads using Cutadapt to remove adapter sequences^66^. The processed reads were collapsed using FASTX Collapser to group identical sequences and reduce PCR-introduced redundancy. Following this, random 4-nucleotide sequences, unique molecular identifiers (UMIs), at both the 5′ and 3′ ends are trimmed to ensure accurate identification of miRNAs. Differentially expressed miRNAs were determined via input of raw miRNA counts into DEseq2^68^. For miRNA isoform quantification, isoforms were determined by aligning at least 18 nucleotides (nt) of the collapsed reads to a reference miRNA database. The count of such matching reads was used to quantify miRNA expression levels and their isoforms.

#### SHAPE-MaP analysis

PacBio *Kah*-specific SHAPE libraries were demultiplexed from one another and converted from .bam to .fastq files using isoseq3. PCR duplicates were collapsed based on a 6 nucleotide UMI included in the *Kah*-specific SHAPE RT primer as described previously. Deduplicated reads were used as inputs for SHAPE-MaP analysis with SHAPEmapper2 (v2.3)^51^ referencing the *Kah* sequence amplified. Nucleotide SHAPE reactivities were calculated via SHAPEmapper2, with nucleotides not mapping to the reference being set to 0. SHAPEmapper2-generated consensus structures (.ct) were used as the reference for structure generation in RNAcanvas/RNA2drawer^69^. SHAPE reactivities <0.5 were considered non-reactive, 0.5-1 moderately reactive, and >1 highly reactive. All sequencing data for this study have been uploaded to the NCBI Sequence Read Archive (SRA) database under SRA accession number PRJNA1189499.

## Acknowledgments

We would like to first sincerely thank our lab members for their thoughtful advice and guidance during the course of this study. We would like to thank David Bartel for sharing S2 cell lines, without which this study would not have been possible. We would like to thank Narry Kim for sharing their detailed AQ-seq protocol used several times in this study. In terms of outside support, we would like to thank Jodi Bubenik for their thoughtful discussions about troubleshooting long-read SHAPE library construction. We would also like to acknowledge the exceptional work done for this study that is still ongoing with our collaborator Jialu Liang and Dr. Qianqian Song. The support from the above-mentioned parties was integral to the success of this study and all should be commended. This work is supported by grants from the National Institutes of Health (R35GM128753 and R01CA282812 to M.X. and T32AI007110 to C.M.T.), the American Cancer Society (Research Scholar Award RSG-21-118-01-RMC to M.X.).

## Author contributions

The study was conceptualized by N.M.H and M.X.; all experiments were performed by N.M.H. with assistance from P.S., T.L., M.Y., and Y.W.; bioinformatics analyses were performed by L.L., N.M.H. and C.M.T.; writing – original draft N.M.H. and M.X.; writing – review & editing L.L., N.M.H., C.M.T., and M.X.; supervision, M.X.; funding for this study was acquired by N.M.H. and M.X.

## REFERENCES

1. Bartel, D.P. (2018). Metazoan MicroRNAs. Cell 173, 20–51. 10.1016/j.cell.2018.03.006.

2. Lee, R.C., Feinbaum, R.L., and Ambrost, V. (1993). The C. elegans Heterochronic Gene lin-4 Encodes Small RNAs with Antisense Complementarity to lin-14. Cell 75, 843–854. 10.1016/0092-8674(93)90529-y

3. Wightman, B., Ha, L., and Ruvkun, G. (1993). Posttranscriptional Regulation of the Heterochronic Gene lin-14 by W-4 Mediates Temporal Pattern Formation in C. elegans. Cell 75, 855–862. 10.1016/0092-8674(93)90530-4

4. Friedman, R.C., Farh, K.K.H., Burge, C.B., and Bartel, D.P. (2009). Most mammalian mRNAs are conserved targets of microRNAs. Genome Res 19, 92–105. 10.1101/gr.082701.108.

5. Agarwal, V., Bell, G.W., Nam, J.-W., and Bartel, D.P. (2015). Predicting effective microRNA target sites in mammalian mRNAs. Elife 4, e05005. 10.7554/eLife.05005.

6. Reinhart, B.J., Slack, F.J., Basson, M., Pasquinelli, A.E., Bettinger, J.C., Rougvie, A.E., Horvitz, H.R., and Ruvkun, G. (2000). The 21-nucleotide let-7 RNA regulates developmental timing in Caenorhabditis elegans. Nature 403, 901–906. 10.1038/35002607.

7. Brennecke, J., Hipfner, D.R., Stark, A., Russell, R.B., and Cohen, S.M. (2003). bantam Encodes a Developmentally Regulated microRNA that Controls Cell Proliferation and Regulates the Proapoptotic Gene hid in Drosophila. Cell 113, 25–36. 10.1016/S0092-8674(03)00231-9.

8. Treiber, T., Treiber, N., and Meister, G. (2019). Regulation of microRNA biogenesis and its crosstalk with other cellular pathways. Nat Rev Mol Cell Biol 20, 5–20. 10.1038/s41580-018-0059-1.

9. Cazalla, D., Yario, T., and Steitz, J. (2010). Down-regulation of a host MicroRNA by a herpesvirus saimiri noncoding RNA. Science (1979) 328, 1563–1566. 10.1126/science.1187197.

10. Ameres, S., Horwich, M., and Zamore, P. (2010). Target RNA–Directed Trimming andTailing of Small Silencing RNAs. Science (1979) 34, 1534–1540.

11. Bitetti, A., Mallory, A.C., Golini, E., Carrieri, C., Carreño Gutiérrez, H., Perlas, E., Pérez-Rico, Y.A., Tocchini-Valentini, G.P., Enright, A.J., Norton, W.H.J., et al. (2018). MicroRNA degradation by a conserved target RNA regulates animal behavior. Nat Struct Mol Biol 25, 244–251. 10.1038/s41594-018-0032-x.

12. Kleaveland, B., Shi, C.Y., Stefano, J., and Bartel, D.P. (2018). A Network of Noncoding Regulatory RNAs Acts in the Mammalian Brain. Cell 174, 350–362.e17. 10.1016/j.cell.2018.05.022.

13. 13. de la Mata, M., Gaidatzis, D., Vitanescu, M., Stadler, M.B., Wentzel, C., Scheiffele, P., Filipowicz, W., and Großhans, H. (2015). Potent degradation of neuronal miRNAs induced by highly complementary targets. EMBO Rep 16, 500–511–511. 10.15252/embr.201540078.

14. Libri, V., Helwak, A., Miesen, P., Santhakumar, D., Borger, J.G., Kudla, G., Grey, F., Tollervey, D., and Buck, A.H. (2012). Murine cytomegalovirus encodes a miR-27 inhibitor disguised as a target. Proc Natl Acad Sci U S A 109, 279–284. 10.1073/pnas.1114204109.

15. Lee, S., Song, J., Kim, S., Kim, J., Hong, Y., Kim, Y., Kim, D., Baek, D., and Ahn, K. (2013). Selective degradation of host MicroRNAs by an intergenic HCMV noncoding RNA accelerates virus production. Cell Host Microbe 13, 678–690. 10.1016/j.chom.2013.05.007.

16. Han, J., Lavigne, C.A., Jones, B.T., Zhang, H., Gillett, F., and Mendell, J.T. (2020). A ubiquitin ligase mediates target-directed microRNA decay independently of tailing and trimming. Science (1979) 370. 10.1126/science.abc9546.

17. Shi, C.Y., Kingston, E.R., Kleaveland, B., Lin, D.H., Stubna, M.W., and Bartel, D.P. (2020). The ZSWIM8 ubiquitin ligase mediates target-directed microRNA degradation. Science (1979) 370. 10.1126/science.abc9359.

18. Han, J., and Mendell, J.T. (2023). MicroRNA turnover: a tale of tailing, trimming, and targets. Trends Biochem Sci 48, 26–39. 10.1016/j.tibs.2022.06.005.

19. Sheu-Gruttadauria, J., Pawlica, P., Klum, S.M., Wang, S., Yario, T.A., Schirle Oakdale, N.T., Steitz, J.A., and MacRae, I.J. (2019). Structural Basis for Target-Directed MicroRNA Degradation. Mol Cell 75, 1243–1255.e7. 10.1016/j.molcel.2019.06.019.

20. Donnelly, B.F., Yang, B., Grimme, A.L., Vieux, K.F., Liu, C.Y., Zhou, L., and McJunkin, K. (2022). The developmentally timed decay of an essential microRNA family is seed-sequence dependent. Cell Rep 40. 10.1016/j.celrep.2022.111154.

21. Kingston, E.R., Blodgett, L.W., and Bartel, D.P. (2022). Endogenous transcripts direct microRNA degradation in Drosophila, and this targeted degradation is required for proper embryonic development. Mol Cell 82, 3872–3884.e9. 10.1016/j.molcel.2022.08.029.

22. Stubna, M.W., Shukla, A., and Bartel, D.P. (2024). Widespread destabilization of C. elegans microRNAs by the E3 ubiquitin ligase EBAX-1. RNA. 10.1261/rna.080276.124.

23. Kingston, E.R., and Bartel, D.P. (2021). Ago2 protects Drosophila siRNAs and microRNAs from target-directed degradation, even in the absence of 2′-O-methylation. 10.1261/rna.

24. Jones, B.T., Han, J., Zhang, H., Hammer, R.E., Evers, B.M., Rakheja, D., Acharya, A., and Mendell, J.T. (2023). Target-directed microRNA degradation regulates developmental microRNA expression and embryonic growth in mammals. Genes Dev 37, 661–674. 10.1101/gad.350906.123.

25. Shi, C.Y., Elcavage, L.E., Chivukula, R.R., Stefano, J., Kleaveland, B., and Bartel, D.P. (2023). ZSWIM8 destabilizes many murine microRNAs and is required for proper embryonic growth and development Running title: Influence of ZSWIM8 on miRNAs in mouse development. Genome research, 33(9), 1482–1496. 10.1101/gr.278073.123

26. Hiers, N.M., Li, T., Traugot, C.M., and Xie, M. (2024). Target-directed microRNA degradation: Mechanisms, significance, and functional implications. WIREs RNA 15, e1832. 10.1002/wrna.1832.

27. Ghini, F., Rubolino, C., Climent, M., Simeone, I., Marzi, M.J., and Nicassio, F. (2018). Endogenous transcripts control miRNA levels and activity in mammalian cells by target-directed miRNA degradation. Nat Commun 9. 10.1038/s41467-018-05182-9.

28. Simeone, I., Rubolino, C., Noviello, T.M.R., Farinello, D., Cerulo, L., Marzi, M.J., and Nicassio, F. (2022). Prediction and pan-cancer analysis of mammalian transcripts involved in target directed miRNA degradation. Nucleic Acids Res 50, 2019–2035. 10.1093/nar/gkac057.

29. Li, L., Sheng, P., Li, T., Fields, C.J., Hiers, N.M., Wang, Y., Li, J., Guardia, C.M., Licht, J.D., and Xie, M. (2021). Widespread microRNA degradation elements in target mRNAs can assist the encoded proteins. Genes Dev 35, 1595–1609. 10.1101/gad.348874.121.

30. Sheng, P., Li, L., Li, T., Wang, Y., Hiers, N.M., Mejia, J.S., Sanchez, J.S., Zhou, L., and Xie, M. (2023). Screening of Drosophila microRNA-degradation sequences reveals Argonaute1 mRNA’s role in regulating miR-999. Nat Commun 14, 2108. 10.1038/s41467-023-37819-9.

31. Buhagiar, A.F., and Kleaveland, B. (2024). To kill a microRNA: emerging concepts in target-directed microRNA degradation. Nucleic Acids Res 52, 1558–1574. 10.1093/nar/gkae003.

32. 32. Yang, A., Bofill-De Ros, X., Stanton, R., Shao, T.J., Villanueva, P., and Gu, S. (2022). TENT2, TUT4, and TUT7 selectively regulate miRNA sequence and abundance. Nat Commun 13, 1–15. 10.1038/s41467-022-32969-8.

33. 33. Yang, A., Shao, T.J., Bofill-De Ros, X., Lian, C., Villanueva, P., Dai, L., and Gu, S. (2020). AGO-bound mature miRNAs are oligouridylated by TUTs and subsequently degraded by DIS3L2. Nat Commun 11, 1–13. 10.1038/s41467-020-16533-w.

34. Helwak, A., and Tollervey, D. (2014). Mapping the miRNA interactome by cross-linking ligation and sequencing of hybrids (CLASH). Nat Protoc 9, 711–728. 10.1038/nprot.2014.043.

35. Moore, M.J., Scheel, T.K.H., Luna, J.M., Park, C.Y., Fak, J.J., Nishiuchi, E., Rice, C.M., and Darnell, R.B. (2015). miRNA–target chimeras reveal miRNA 3′-end pairing as a major determinant of Argonaute target specificity. Nat Commun 6, 8864. 10.1038/ncomms9864.

36. 36. Manakov, S.A., Shishkin, A.A., Yee, B.A., Shen, K.A., Cox, D.C., Park, S.S., Foster, H.M., Chapman, K.B., Yeo, G.W., and Van Nostrand, E.L. (2022). Scalable and deep profiling of mRNA targets for individual microRNAs with chimeric eCLIP. bioRxiv, 2022.02.13.480296. 10.1101/2022.02.13.480296.

37. A, G.L., Sunantha, S., Merin, T., C, T.P., and Rolf, R. (2018). Modified Cross-Linking, Ligation, and Sequencing of Hybrids (qCLASH) Identifies Kaposi’s Sarcoma-Associated Herpesvirus MicroRNA Targets in Endothelial Cells. J Virol 92, 10.1128/jvi.02138-17. 10.1128/jvi.02138-17.

38. Kim, H., Kim, J., Kim, K., Chang, H., You, K. V., and Kim, N. (2019). Bias-minimized quantification of microRNA reveals widespread alternative processing and 3 end modification. Nucleic Acids Res 47, 2630–2640. 10.1093/nar/gky1293.

39. Jayaprakash, A.D., Jabado, O., Brown, B.D., and Sachidanandam, R. (2011). Identification and remediation of biases in the activity of RNA ligases in small-RNA deep sequencing. Nucleic Acids Res 39, e141–e141. 10.1093/nar/gkr693.

40. Zhuang, F., Fuchs, R.T., Sun, Z., Zheng, Y., and Robb, G.B. (2012). Structural bias in T4 RNA ligase-mediated 3′-adapter ligation. Nucleic Acids Res 40, e54–e54. 10.1093/nar/gkr1263.

41. Hafner, M., Renwick, N., Brown, M., Mihailović, A., Holoch, D., Lin, C., Pena, J.T.G., Nusbaum, J.D., Morozov, P., Ludwig, J., et al. (2011). RNA-ligase-dependent biases in miRNA representation in deep-sequenced small RNA cDNA libraries. RNA 17, 1697– 1712. 10.1261/rna.2799511.

42. Miller, B.R., Wei, T., Fields, C.J., Sheng, P., and Xie, M. (2018). Near-infrared fluorescent northern blot. Rna 24, 1871–1877. 10.1261/rna.068213.118.

43. Fang, W., and Bartel, D.P. (2020). MicroRNA Clustering Assists Processing of Suboptimal MicroRNA Hairpins through the Action of the ERH Protein. Mol Cell 78, 289–302.e6. 10.1016/j.molcel.2020.01.026.

44. Shang, R., Baek, S.C., Kim, K., Kim, B., Kim, V.N., and Lai, E.C. (2020). Genomic Clustering Facilitates Nuclear Processing of Suboptimal Pri-miRNA Loci. Mol Cell 78, 303–316.e4. 10.1016/j.molcel.2020.02.009.

45. Agarwal, V., Subtelny, A.O., Thiru, P., Ulitsky, I., and Bartel, D.P. (2018). Predicting microRNA targeting efficacy in Drosophila. Genome Biol 19, 152. 10.1186/s13059-018-1504-3.

46. 46. Corey, D.R., and Abrams, J.M. (2001). Morpholino antisense oligonucleotides: tools for investigating vertebrate development. Genome Biol 2, reviews1015.1. 10.1186/gb-2001-2-5-reviews1015.

47. Staton, A.A., and Giraldez, A.J. (2011). Use of target protector morpholinos to analyze the physiological roles of specific miRNA-mRNA pairs in vivo. Nat Protoc 6, 2035–2049. 10.1038/nprot.2011.423.

48. Siegfried, N.A., Busan, S., Rice, G.M., Nelson, J.A.E., and Weeks, K.M. (2014). RNA motif discovery by SHAPE and mutational profiling (SHAPE-MaP). Nat Methods 11, 959–965. 10.1038/nmeth.3029.

49. Smola, M.J., Rice, G.M., Busan, S., Siegfried, N.A., and Weeks, K.M. (2015). Selective 2′-hydroxyl acylation analyzed by primer extension and mutational profiling (SHAPE-MaP) for direct, versatile and accurate RNA structure analysis. Nat Protoc 10, 1643–1669. 10.1038/nprot.2015.103.

50. Spitale, R.C., Crisalli, P., Flynn, R.A., Torre, E.A., Kool, E.T., and Chang, H.Y. (2013). RNA SHAPE analysis in living cells. Nat Chem Biol 9, 18–20. 10.1038/nchembio.1131.

51. Busan, S., and Weeks, K.M. (2018). Accurate detection of chemical modifications in RNA by mutational profiling (MaP) with ShapeMapper 2. RNA 24, 143–148. 10.1261/rna.061945.117.

52. Jones, A.N., Pisignano, G., Pavelitz, T., White, J., Kinisu, M., Forino, N., Albin, D., and Varani, G. (2020). An evolutionarily conserved RNA structure in the functional core of the lincRNA Cyrano. RNA 26, 1234–1246. 10.1261/rna.076117.120.

53. Guo, L.-T., Adams, R.L., Wan, H., Huston, N.C., Potapova, O., Olson, S., Gallardo, C.M., Graveley, B.R., Torbett, B.E., and Pyle, A.M. (2020). Sequencing and Structure Probing of Long RNAs Using MarathonRT: A Next-Generation Reverse Transcriptase. J Mol Biol 432, 3338–3352. 10.1016/j.jmb.2020.03.022.

54. Schirle, N.T., Sheu-Gruttadauria, J., and MacRae, I.J. (2014). Structural basis for microRNA targeting. Science (1979) 346, 608–613. 10.1126/science.1258040.

55. Chandradoss, S.D., Schirle, N.T., Szczepaniak, M., MacRae, I.J., and Joo, C. (2015). A Dynamic Search Process Underlies MicroRNA Targeting. Cell 162, 96–107. 10.1016/j.cell.2015.06.032.

56. McGeary, S.E., Lin, K.S., Shi, C.Y., Pham, T.M., Bisaria, N., Kelley, G.M., and Bartel, D.P. (2019). The biochemical basis of microRNA targeting efficacy. Science (1979) 366, eaav1741. 10.1126/science.aav1741.

57. Kosek, D.M., Banijamali, E., Becker, W., Petzold, K., and Andersson, E.R. (2023). Efficient 3′-pairing renders microRNA targeting less sensitive to mRNA seed accessibility. Nucleic Acids Res 51, 11162–11177. 10.1093/nar/gkad795.

58. Broughton, J.P., Lovci, M.T., Huang, J.L., Yeo, G.W., and Pasquinelli, A.E. (2016). Pairing beyond the Seed Supports MicroRNA Targeting Specificity. Mol Cell 64, 320–333. 10.1016/j.molcel.2016.09.004.

59. Salomon, W.E., Jolly, S.M., Moore, M.J., Zamore, P.D., and Serebrov, V. (2015). Single-Molecule Imaging Reveals that Argonaute Reshapes the Binding Properties of Its Nucleic Acid Guides. Cell 162, 84–95. 10.1016/j.cell.2015.06.029.

60. Wee, L.M., Flores-Jasso, C.F., Salomon, W.E., and Zamore, P.D. (2012). Argonaute Divides Its RNA Guide into Domains with Distinct Functions and RNA-Binding Properties. Cell 151, 1055–1067. 10.1016/j.cell.2012.10.036.

61. Mendoza-Garcia, P., Basu, S., Sukumar, S.K., Arefin, B., Wolfstetter, G., Anthonydhason, V., Molander, L., Uçkun, E., Lindehell, H., Lebrero-Fernandez, C., et al. (2021). DamID transcriptional profiling identifies the Snail/Scratch transcription factor Kahuli as an Alk target in the Drosophila visceral mesoderm. Development 148, dev199465. 10.1242/dev.199465.

62. Cayirlioglu, P., Kadow, I.G., Zhan, X., Okamura, K., Suh, G.S.B., Gunning, D., Lai, E.C., and Zipursky, S.L. (2008). Hybrid Neurons in a MicroRNA Mutant Are Putative Evolutionary Intermediates in Insect CO2 Sensory Systems. Science (1979) 319, 1256–1260. 10.1126/science.1149483.

63. 63. Sun, K., Jee, D., de Navas, L.F., Duan, H., and Lai, E.C. (2015). Multiple In Vivo Biological Processes Are Mediated by Functionally Redundant Activities of Drosophila mir-279 and mir-996. PLoS genetics, 11(6), e1005245. 10.1371/journal.pgen.1005

64. 64. Duan, H., de Navas, L.F., Hu, F., Sun, K., Mavromatakis, Y.E., Viets, K., Zhou, C., Kavaler, J., Johnston, R.J., Tomlinson, A., et al. (2018). The mir-279/996 cluster represses receptor tyrosine kinase signaling to determine cell fates in the Drosophila eye. Development 145, dev159053. 10.1242/dev.159053.

65. Rajman, M., and Schratt, G. (2017). MicroRNAs in neural development: from master regulators to fine-tuners. Development 144, 2310–2322. 10.1242/dev.144337.

66. Martin, M. (2011). Cutadapt removes adapter sequences from high-throughput sequencing reads. EMBnet J 17, 10–12.

67. Kim, D., Paggi, J.M., Park, C., Bennett, C., and Salzberg, S.L. (2019). Graph-based genome alignment and genotyping with HISAT2 and HISAT-genotype. Nat Biotechnol 37, 907–915. 10.1038/s41587-019-0201-4.

68. Love, M.I., Huber, W., and Anders, S. (2014). Moderated estimation of fold change and dispersion for RNA-seq data with DESeq2. Genome Biol 15, 550. 10.1186/s13059-014-0550-8.

69. Johnson, P.Z., Kasprzak, W.K., Shapiro, B.A., and Simon, A.E. (2019). RNA2Drawer: geometrically strict drawing of nucleic acid structures with graphical structure editing and highlighting of complementary subsequences. RNA Biol 16, 1667–1671. 10.1080/15476286.2019.1659081.

